# Gene size matters: What determines gene length in the human genome?

**DOI:** 10.1101/2020.01.10.901272

**Authors:** Inês Lopes, Gulam Altab, Priyanka Raina, João Pedro de Magalhães

## Abstract

While it is expected for gene length to be influenced by factors such as intron number and evolutionary conservation, we have yet to fully understand the connection between gene length and function in the human genome.

In this study, we show that, as expected, there is a strong positive correlation between gene length and the number of SNPs, introns and protein size. Amongst tissue specific genes, we find that the longest genes are expressed in blood vessels, nerve, thyroid, cervix uteri and brain, while the smallest genes are expressed within the pancreas, skin, stomach, vagina and testis. We report, as shown previously, that natural selection suppresses changes for genes with longer lengths and promotes changes for smaller genes. We also observed that longer genes have a significantly higher number of co-expressed genes and protein-protein interactions. In the functional analysis, we show that bigger genes are often associated with neuronal development, while smaller genes tend to play roles in skin development and in the immune system. Furthermore, pathways related to cancer, neurons and heart diseases tend to have longer genes, with smaller genes being present in pathways related to immune response and neurodegenerative diseases.

We hypothesise that longer genes tend to be associated with functions that are important early in life, while smaller genes play a role in functions that are important throughout the organisms’ whole life, like the immune system which require fast responses.

**Author Summary:** Even though the human genome has been fully sequenced, we still do not fully grasp all of its nuances. One such nuance is the length of the genes themselves. Why are certain genes longer than others? Is there a common function shared by longer/smaller genes? What exactly makes gene longer? We tried answering these questions using a variety of analysis. We found that, while there was not a particular strong factor in genes that influenced their size, there could be an influence of several gene characteristics in determining the length of a gene. We also found that longer genes are linked with the development of neurons, cancer, heart diseases and muscle cells, while smaller genes seem to be mostly related with the immune system and the development of the skin. This led us to believe that, whether the gene has an important function early in our life, or throughout our whole lives, or even if the function requires a rapid response, that its gene size will be influenced accordingly.

## Background

With the sequencing of the human genome [1–3] there arose a great interest in understanding the relationship between genotype and phenotype, especially concerning human health [4, 5]. However, despite the recent advancements, we have yet to fully understand the human genome and its complexity [6].

Several studies have tried to decipher a connection between the length of a gene and its function. It is believed that genes that are more evolutionarily conserved are often associated with longer gene length and higher intronic burden [7–10]. In contrast, smaller gene length is often associated with high expression, smaller proteins and little intronic content [11]. This hypothesis is further supported by the house keeping genes, which are widely expressed and have characteristics similar to smaller gene length genes [12]. It was hypothesised that, due to this great levels of expression for smaller genes, there is selective pressure to maximize protein synthesis efficiency [11]. If that is the case, then the next question should be what functions serve longer genes to compensate for their expensive production of proteins. Gene length has been importantly associated with biological timing. The smaller genes produce smaller proteins faster, and these proteins often play a part in the regulation of longer proteins, which are expressed much later into the response. This allows for regulatory mechanisms to be set up in preparation for important protein expression [13]. On the other hand, longer genes have been associated with some important processes, including embryonic development [14] and neuronal processes [15]. Longer genes have also been previously shown to be related to diseases such as cancer, cardiomyopathies and diabetes [15].

In this present work, we used human genome data [16], to identify possible functions based on gene size. Correlation tests were used to search for relationships between gene length and other gene characteristics. In order to find the specific functions associated with gene size, the Gene Ontology (GO) and the KEGG Pathway were used. We observed that longer genes are expressed in the brain, heart diseases and cancer, while smaller genes mostly participate in the immune system and in the development of the skin. Therefore, we hypothesize that genes with longer lengths are mostly associated with functions in the early development stages, while genes with smaller lengths have important roles in day-to-day functions.

## Results

### Longest and shortest genes

For all of the protein-coding transcripts in the human genome, a dataset was built selecting only the transcripts with the highest transcript length per gene (N=19,714 genes, S1 Table). Using mostly the transcript length for the rest of this analysis, stems from the fact that there is a very high correlation between the length of the longest transcript of a gene and its respective gene length (S1 Fig, Kendall test, tau = 0.72, p-value < 2.20E-16). The 5 biggest genes in terms of transcript length have all been studied previously, and we can see that they are associated with neuron functions [17–19], cardiac tissue [20] and cancer [21] (Table 1). However, the smallest genes might be annotation errors in the genome build.

**Table 1.**
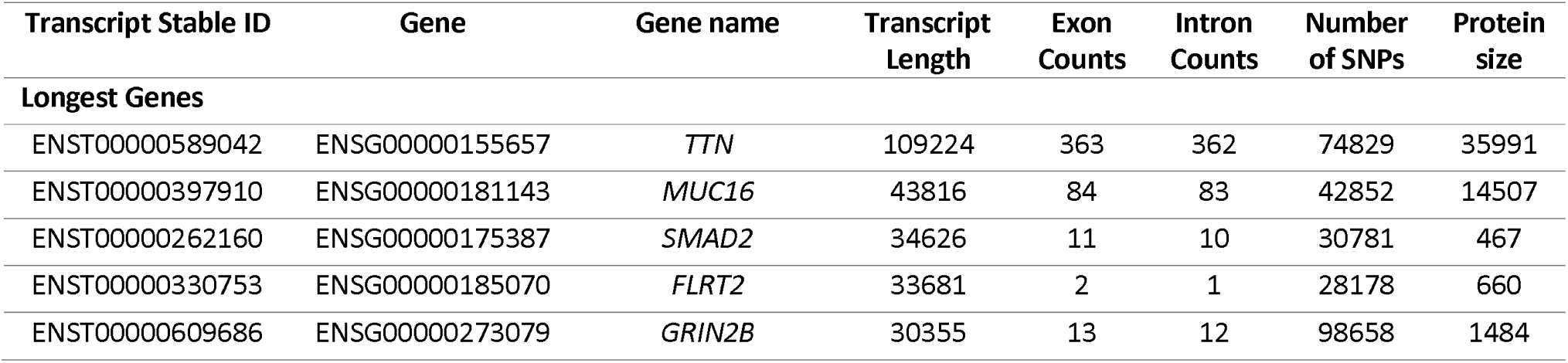
List of the top 5 longest protein-coding transcripts in human.

| Transcript Stable ID | Gene | Gene name | Transcript Length | Exon Counts | Intron Counts | Number of SNPs | Protein size |
| --- | --- | --- | --- | --- | --- | --- | --- |
| **Longest Genes** |  |  |  |  |  |  |  |
| ENST00000589042 | ENSG00000155657 | *TTN* | 109224 | 363 | 362 | 74829 | 35991 |
| ENST00000397910 | ENSG00000181143 | *MUC16* | 43816 | 84 | 83 | 42852 | 14507 |
| ENST00000262160 | ENSG00000175387 | *SMAD2* | 34626 | 11 | 10 | 30781 | 467 |
| ENST00000330753 | ENSG00000185070 | *FLRT2* | 33681 | 2 | 1 | 28178 | 660 |
| ENST00000609686 | ENSG00000273079 | *GRIN2B* | 30355 | 13 | 12 | 98658 | 1484 |

### Functional analysis

One of the main objectives of the present study was to understand if gene function changed depending on the gene length. Keeping this in mind, and using a list of the top 5% protein coding genes with the longest and smallest transcript length, we performed an analysis, using tools like WebGestalt [22], DAVID [23, 24], KEGG [25] and Molecular Signature Database [26, 27]. The results for KEGG Pathways, were colour coded for each boxplot based on their association with the terms we found most relevant (brain, cancer, heart, immune system, muscle, neurodegenerative disease, skin and other). For cases where there was no direct association, a literature search was done for relevant articles that might show that genes in those pathways were related to brain [28–47], cancer [48], immune system [49–53] and skin [54–58].

For genes with longer gene length (Fig 1), most of the biological functions found seem to be associated with the brain, specifically in regards to neurons. This can also be confirmed when looking at the Cellular Component (S2A Fig) and Molecular Function (S2B Figure), and at the similar results produced using DAVID (S2 Table).

**Fig 1.**
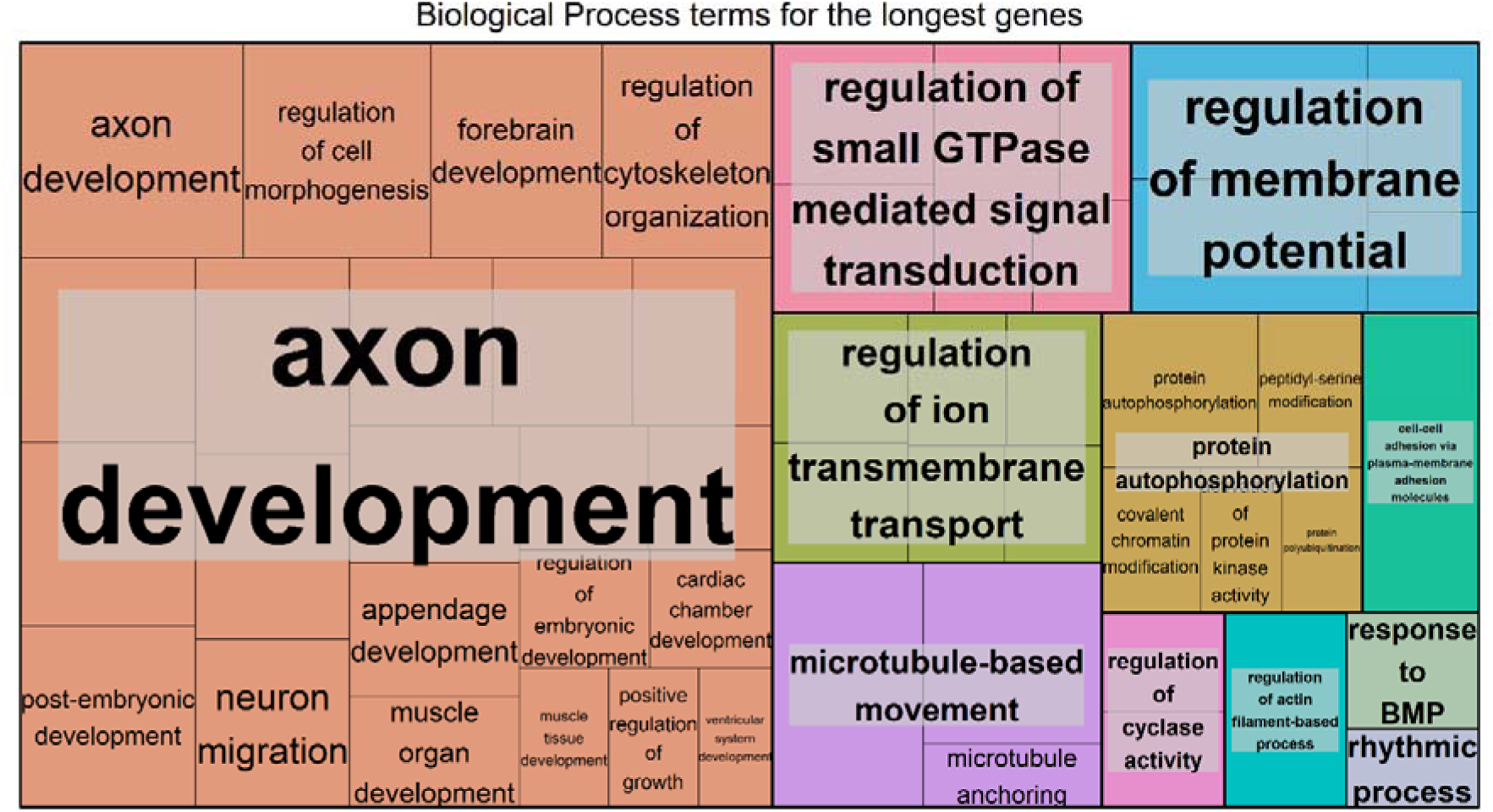
Biological Process terms found associated to genes with the longest transcript length. Overrepresentation Enrichment Analysis was performed with WebGestalt [22] and the visualization tool REViGO [59] was used to produce this figure. The significance level was p<0.05 and the FDR was set at 0.05. FDR estimation was done using the Benjamini–Hochberg method.

For the genes with smaller gene length (Fig 2), most of the biological functions found are related to skin and the immune system. Similarly to what we observed before, Cellular Component (S2C Fig), Molecular Function (S2D Fig) and DAVID (S2 Table) results supported this observation.

**Fig 2.**
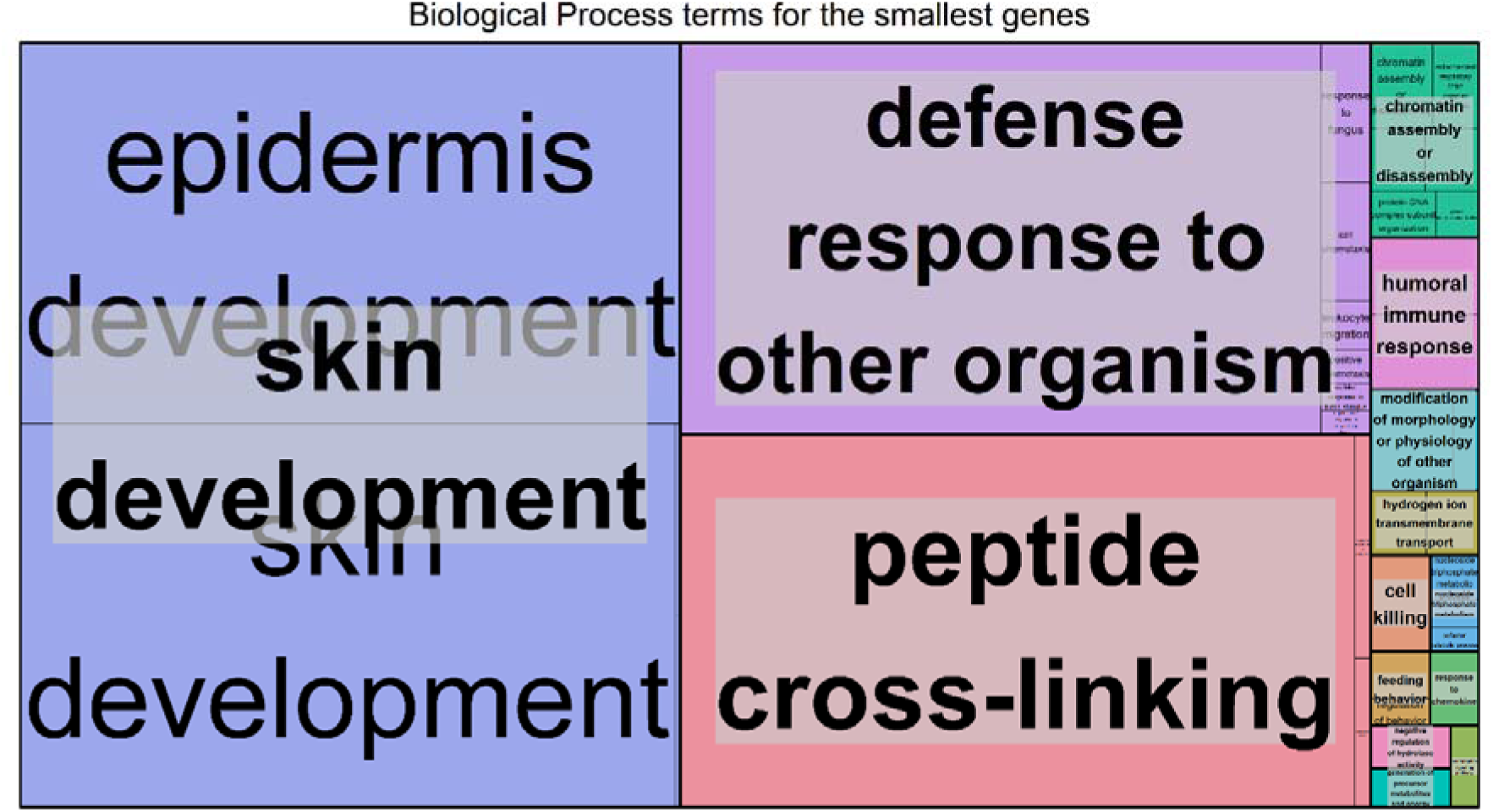
Biological Process terms found associated to genes with the smallest transcript length. Overrepresentation Enrichment Analysis was performed with WebGestalt [22] and the visualization tool REViGO [59] was used to produce this figure. The significance level was p<0.05 and the FDR was set at 0.05. FDR estimation was done using the Benjamini–Hochberg method.

Additionally, while looking at the KEGG Pathways results for longest transcript length, we identified pathways associated with the brain, cancer, heart disease and muscle (Fig 3A, S3 Fig), while the pathways with the smallest transcript length are mostly associated with the immune system, a few of them were also associated with skin and neurodegenerative diseases (Fig 3B, S3 Fig).

**Fig 3.**
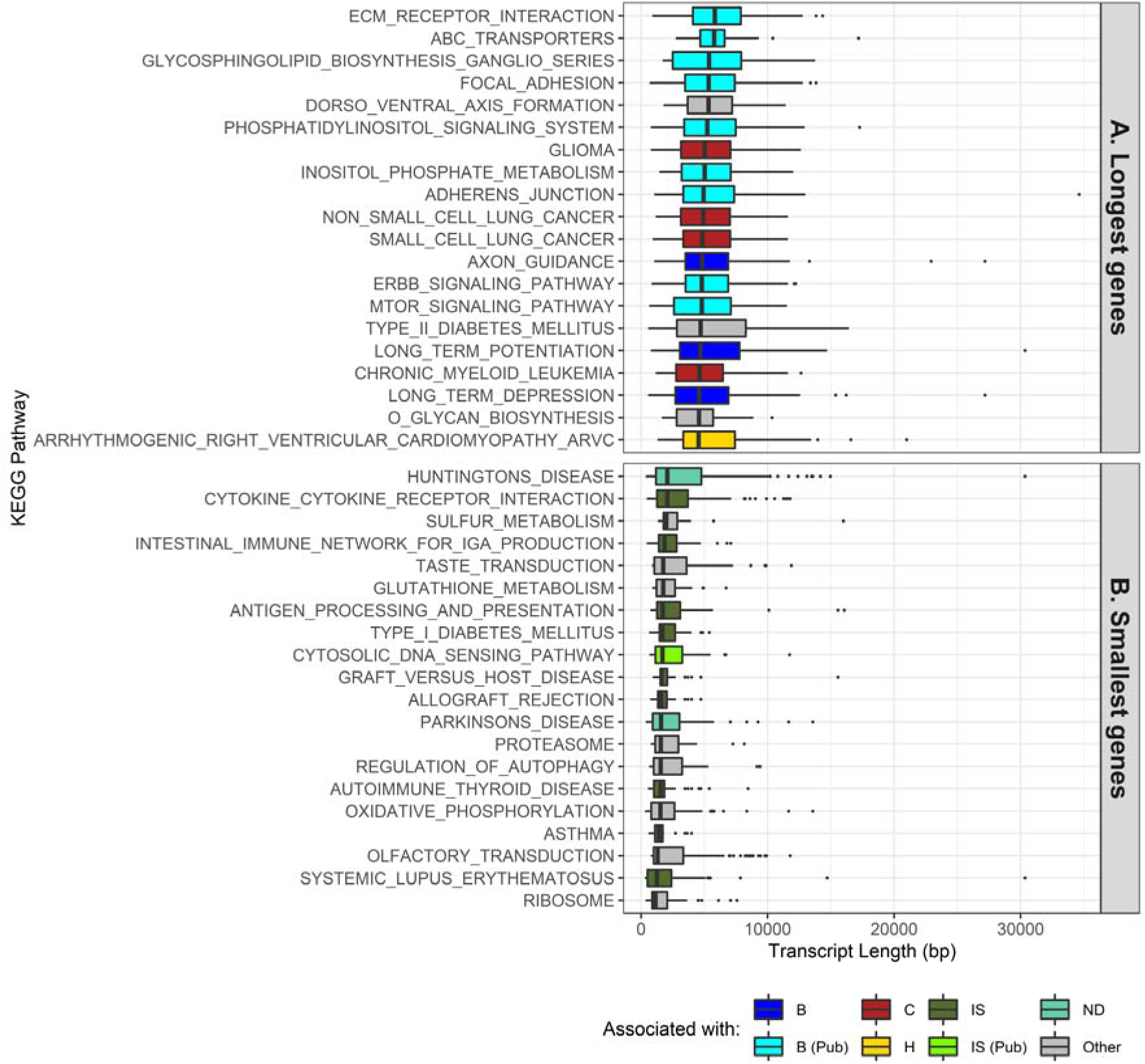
Transcript length distribution per KEGG Pathway for the longest and smallest genes. Colours illustrate what the KEGG pathway has been directly associated with (B for Brain, C for Cancer, H for Heart, IS for Immune system and ND for Neurodegenerative diseases), due to it being stated in the pathway itself, or indirectly associated with (Pub tag), by means of literature references. KEGG Pathways and genes involved in said pathways were obtained from the Molecular Signature Database [26, 27]. <u>A:</u> Top 20 Pathways with the longest genes, ordered by median; <u>B:</u> Top 20 Pathways with the smallest genes, ordered by median.

The full KEGG Results (186 gene sets) can be found in the S3 Fig, and the KEGG Pathway IDs can be found in the S3 Table.

### Gene properties correlate with transcript length

In order to understand the relationship between transcript length and other gene characteristics, a correlation analysis was done. When looking at the number of SNPs for each transcript (Fig 4A), there was a significant positive correlation with transcript length (Kendall test, tau = 0.45, p-value < 2.20E-16). Similar results were found, when comparing the number of SNPs per gene with gene length (S4A Fig, Kendall test, tau = 0.49, p-value < 2.20E-16). After comparing the number of introns and the transcript length (Fig 4B), we found a weak significant positive correlation between these two variables (Kendall test, tau = 0.35, p-value < 2.20E-16). The strongest positive correlation (Kendall test, tau = 0.48, p-value < 2.20E-16) was associated with the protein size (Fig 4C), and the weakest correlation (Kendall test, tau = 0.04, p-value = 3.06E-14) was associated with the average gene expression (Fig 4D).

**Fig 4.**
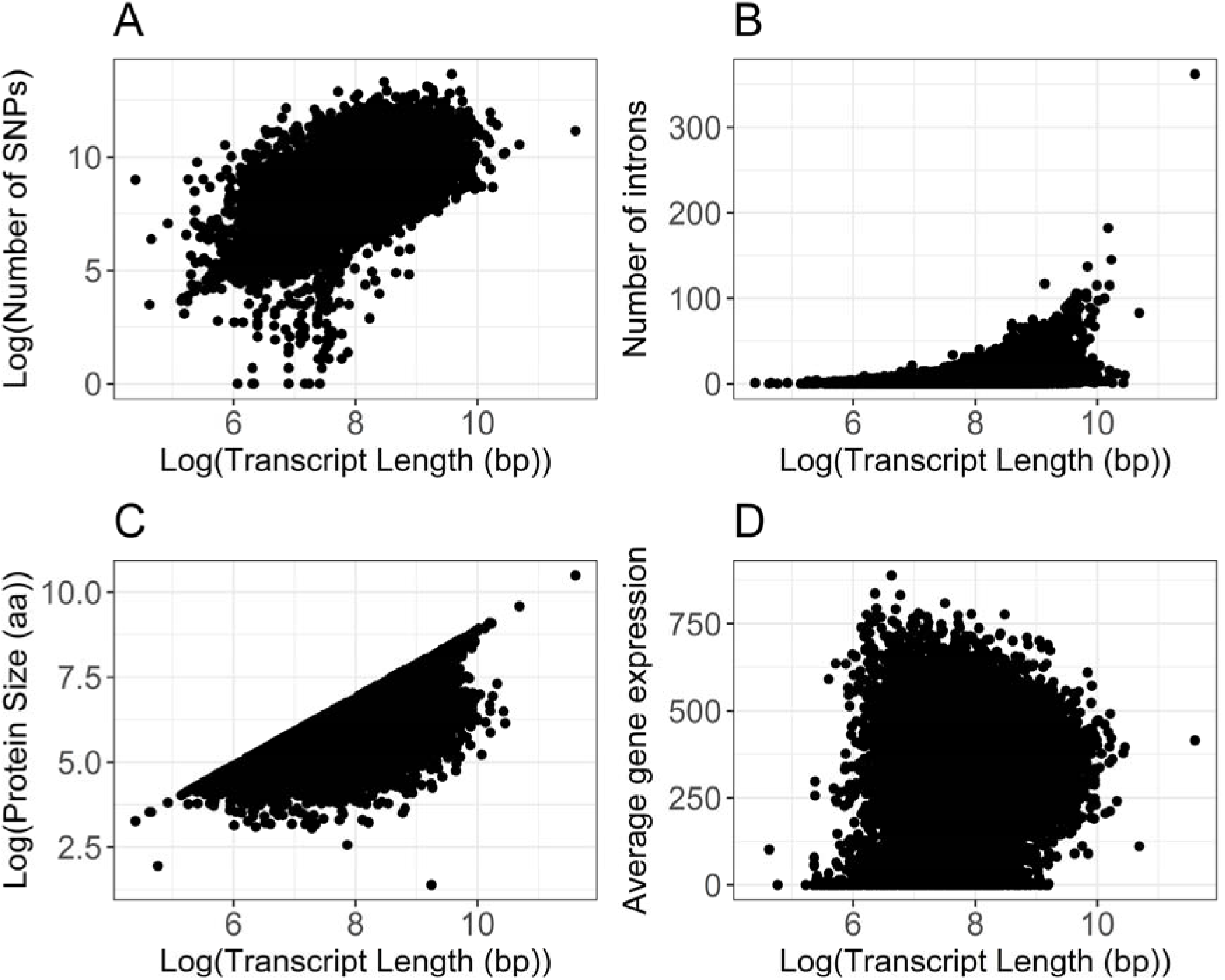
Correlation analysis between Transcript Length (bp) and several other gene characteristics. All figures have been logarithmically transformed in order to help visualize their relationship and/or account for the skewing introduced by outliers. The original versions of the figures can be found in the S4B, S4C, S4D and S4E Fig. <u>A:</u> Correlation between the log transformed number of SNPs and the log transformed Transcript Length (bp) (Kendall test, tau = 0.45, p-value < 2.20E-16). Number of SNPs and Transcript Length for each transcript were obtained using biomart; <u>B:</u> Correlation between the number of introns and the log transformed Transcript Length (bp) (Kendall test, tau = 0.35, p-value < 2.20E-16). Number of introns and Transcript Length for each transcript were obtained using biomart; <u>C:</u> Correlation between the log transformed Protein Size (aa) and the log transformed Transcript Length (bp) (Kendall test, tau = 0.48, p-value < 2.20E-16). Protein Size and Transcript Length were obtained using biomart; <u>D:</u> Correlation between the Average Gene Expression and the log transformed Transcript Length (bp) (Kendall test, tau = 0.04, p-value = 3.06E-14). Average Gene Expression was obtained from the UCSC Genome browser, this value was derived from the total median expression level across all tissues and was based on the GTEx project. Transcript Length was obtained using biomart.

Additionally, for the correlations with Transcript count (S4F Fig) and GC content (S4G Fig), we observed a weak significant positive correlation (Kendall test, tau = 0.22, p-value < 2.20E-16) and a weak significant negative correlation (Kendall test, tau = -0.19, p-value < 2.20E-16), respectively.

We were also interested in understanding the effect of transcript length in some particular mutations. We observed some strong statistically significant correlations between transcript length and synonymous (S4H Fig, Kendall test, tau = 0.44, p-value < 2.20E-16) and missense (S4I Fig, Kendall test, tau = 0.42, p-value < 2.20E-16) mutations. However, in case of nonsense mutations (S4J Fig, Kendall test, tau = 0.21, p-value < 2.20E-16) a weaker significant positive correlation with transcript length was observed. This was followed by the calculation of Missense/Synonymous (MIS/SYN) and Nonsense/Synonymous (NONS/SYN) rates in order to measure the functional importance of gene length. We observed that this ratios had similarly negative correlations with transcript length, with MIS/SYN having a weaker significant correlation (S4K Fig, Kendall test, tau = -0.07, p-value < 2.20E-16) than NONS/SYN (S4L Fig, Kendall test, tau = -0.19, p-value < 2.20E-16).

In order to better understand if the correlations found were solely due to the transcript length or if other factors were influencing them, we built a correlation matrix with several gene characteristics (Fig 5). We observed that properties like intron counts, CDS length, protein size, number of SNPs and transcript count have some strong positive correlations amongst themselves, some of which were stronger than any other correlation with transcript length. This indicated that strong correlations with transcript length might not be due to the sole action of transcript length itself, but rather due to a combined action between several gene characteristics.

**Fig 5.**
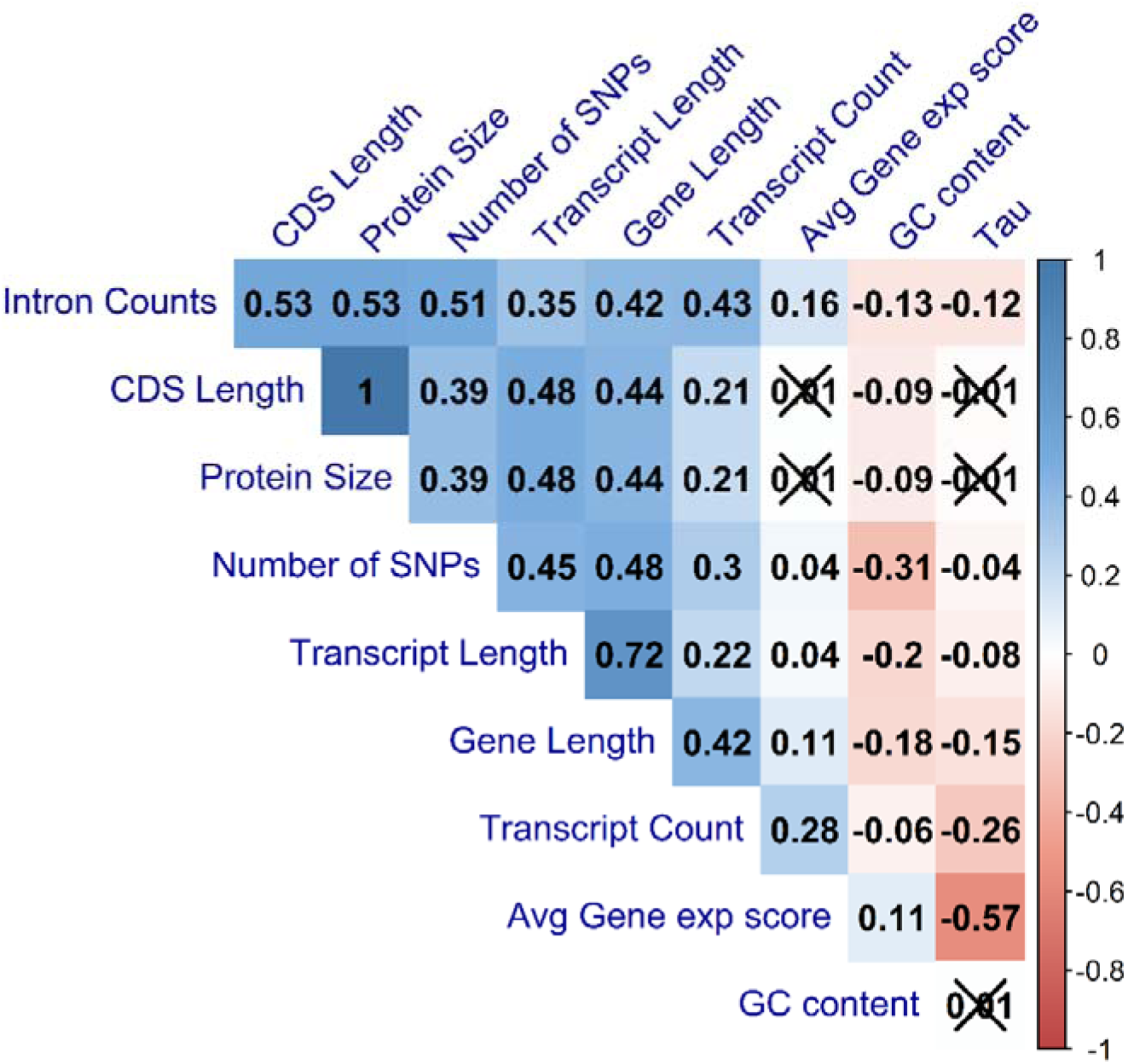
Correlation matrix between gene properties. Kendall’s test was used as a measurement of correlation, with the numbers and the gradient of colours symbolizing the Tau values for each comparison. Number of SNPs values is for each transcript. Values that are crossed out are not statistically significant. Values are clustered together based on their Tau values.

### Distribution of transcript length and expression in human tissues

In this present work we have found that transcript length seems to peak at 2065 bp, with smaller transcripts being more common than longer ones (S5A Fig). As described previously [9], the distribution of the number of introns in the human genome (S5B Fig) has a mode of 3 introns and there are very few genes with a large number of introns. The gene with the most introns is TTN, with 362 introns, which also leads the list of genes with the longest transcript length.

To better understand the distribution of transcript length in the human tissue specific genes, we used Tau values obtained from GTEx data [60]. Tau was used has a measure of tissue specificity, based on the expression profile in different tissues, with values ranging from 0, for broadly expressed genes, to 1, for tissue specific genes [61]. For genes with a Tau value above 0.8 (Fig 6, S6 Fig for the non-log transformed version), we observed that longer tissue specific genes are often associated with the blood vessel, nerve, thyroid, cervix uteri and brain, while smaller tissue specific genes are found in the pancreas, skin, stomach, vagina and testis.

**Fig 6.**
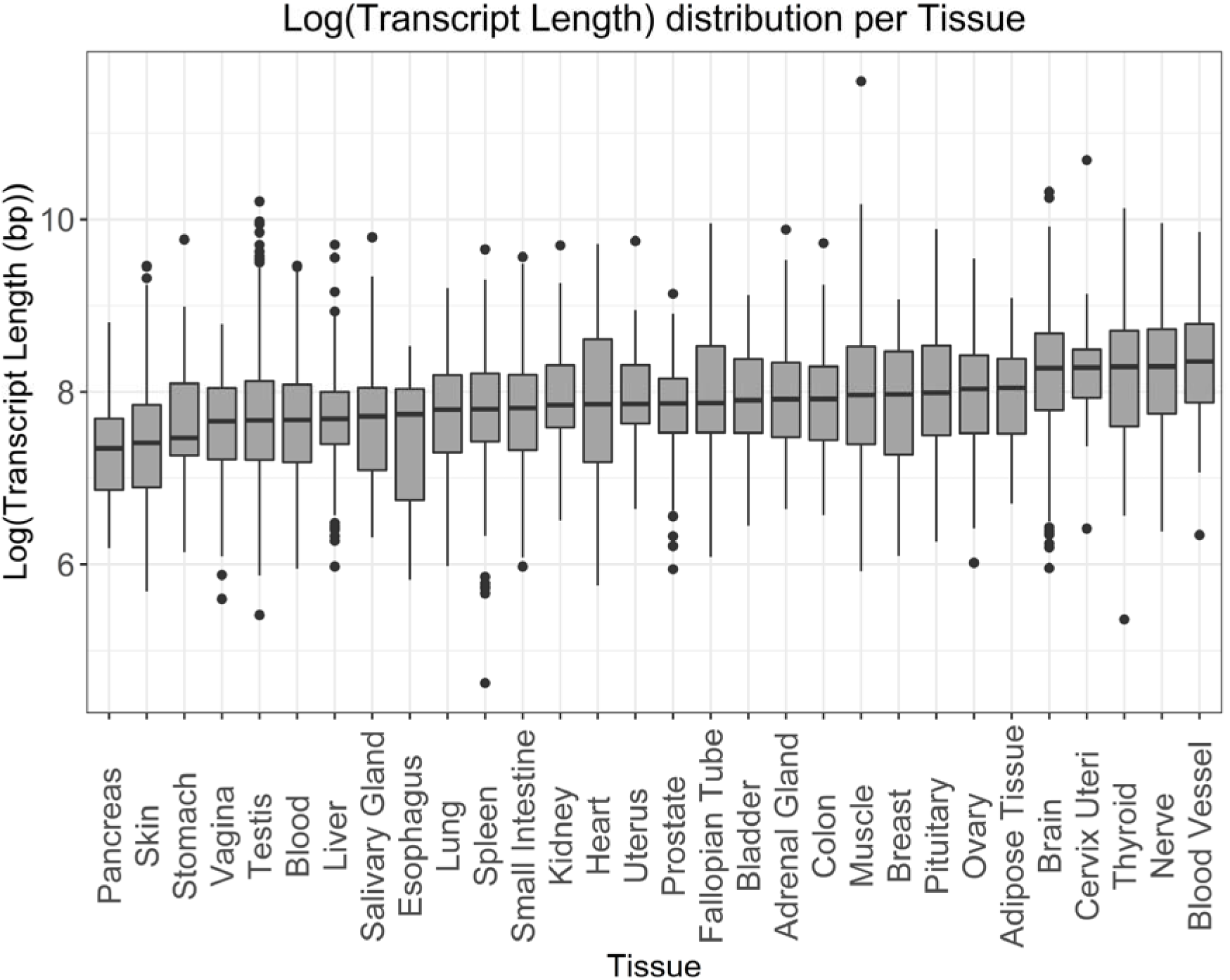
Log transformed Transcript length distribution for genes specifically expressed in the given Tissues. Tissue specificity was defined as a gene having a Tau specificity score greater than 0.8.

**Fig 7.**
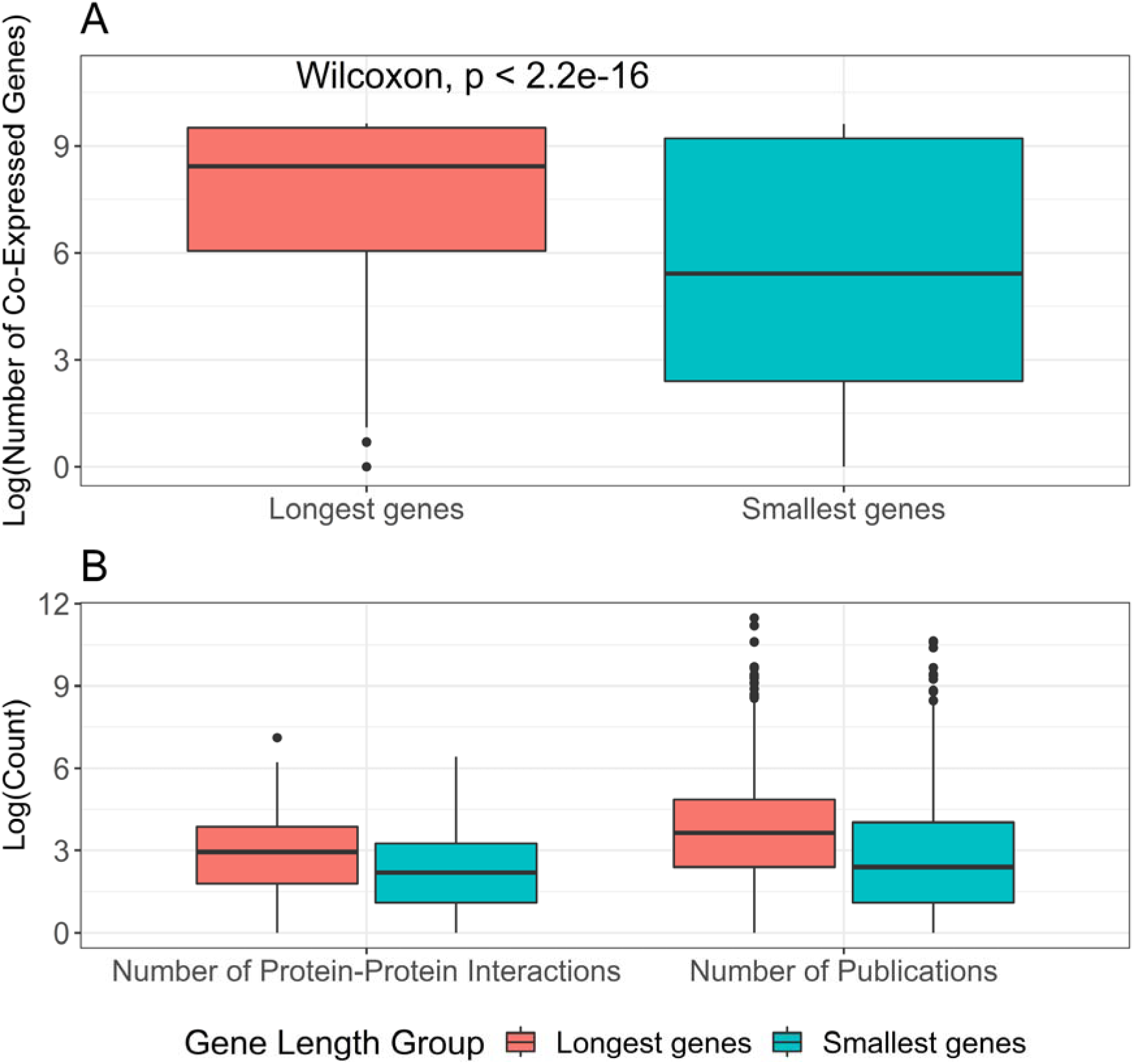
Co-expression and protein-protein Interaction results pertaining to the longest and the smallest genes. The High group corresponds to the top 5% longest genes found in our original dataset (N_High_ = 986), while the Low group corresponds to the top 5% smallest genes found in our original dataset (N_Low_ = 986). <u>A:</u> Distribution of the Log transformed number of co-expressed genes for long genes and small genes. Number of co-expressed genes was obtained from data publicly available in GeneFriends [64]; <u>B:</u> Distribution of the number of protein-protein interactions and the number of publications for longer and smaller genes, all Log transformed. Number of protein-protein interactions was obtained from BioGRID [65] and the number of publications was obtained from PubMed.

### Ageing and transcript length

Ageing is an important factor in our lives, and it affects most organisms. We were curious to see if, for genes related to ageing, the distribution of transcript length was significantly different than the rest of the protein-coding genes. We observed (S7A Fig and S7B Fig) that genes associated with ageing (N = 307) [62] have longer transcript lengths (median = 3517) when compared with the rest of our dataset (median = 2956), and that this difference of medians was significant (Wilcoxon rank sum test, p-value = 0.00036).

To further understand if longer or smaller genes were more prominent with age, we used genes from ageing signatures obtained from a meta-analysis in human, mice and rat [60]. Genes from this signature were either overexpressed (N_Total_ = 449, N_Brain_ = 147, N_Heart_ = 35, N_Muscle_ = 49) or underexpressed (N_Total_ = 162, N_Brain_ = 16, N_Heart_ = 5, N_Muscle_ = 73) with age. Overall, the difference in medians for the distribution of transcript length in genes overexpressed (median = 3068) and underexpressed (median = 3026.5) with ageing was not observed to be significant (S7C Fig, Wilcoxon rank sum test, p-value = 0.81). However, tissue specific signatures showed that the brain favours smaller genes with age (S7D Fig, Wilcoxon rank sum test, p-value = 0.00086, median for overexpression in brain = 2651, median for underexpression in brain = 5824).

### Evolution and transcript length

The relationship between intronic burden and evolution has been established before [9], but very few works approached this on a gene length front. Therefore we obtained the dN and dS values for three organisms paired with human, mouse (S8A Fig), gorilla (S8B Fig) and chimpanzee (S8C Fig), and we aimed to see how the distribution of transcript length happened in function of their dN/dS ratio. Overall, longer genes were associated with a dN/dS ratio lesser to 1 (median transcript length is 3294, 3377 and 3338 for mouse, chimpanzee and gorilla respectively), while smaller genes seem to be more associated with dN/dS ratios above or equal to 1 (median transcript length is 1171.5, 2229.5 and 2092 for mouse, chimpanzee and gorilla respectively) and the median of both groups was always significantly different (Wilcoxon rank sum test, p-value = 0.00073 for mouse and <2.2E-16 for both gorilla and chimpanzee).

### Co-Expression Analysis and Protein-Protein Interactions

Co-expression networks can help us to better understand the functions of genes that are often expressed together [63]. In order to see if the gene length influenced the amount of co-expressed partners, we used data from GeneFriends [64] (S4 Table). We observed a rather weak correlation between transcript length and the number of co-expression partners in our dataset (S9A Fig, Kendall Test, tau = 0.10, p-value < 2.2E-16). However, despite this weak correlation, longer genes appeared to have more co-expressed gene partners than smaller genes (Fig 7A, Wilcoxon rank sum test, p-value < 2.2E-16, not-transformed figure in S9B Fig, median values of co-expression partners for longer genes = 2725, median values of co-expression partners for smaller genes = 32). We further analysed top and lowest hundred human co-expressed genes from the GeneFriends database (S4 Table) and observed that top highly co-expressed genes in the database have significantly higher transcript length (S9C Fig, Wilcoxon rank sum test, p-value = 0.00072, median = 3880) with respect to the bottom ones (median = 2587.5).

To determine if transcript length also influenced the number of protein-protein interactions, we used the protein-protein interaction data from BioGRID [65] (S5 Table). The results obtained were similar to the co-expression, where a weak correlation was observed between transcript length and the number of protein-protein interactions (S10A Fig, Kendall Test, tau = 0.06, p-value < 2.2E-16).

From such results, one would think that publication bias would have an effect on the number of interactions found. So, we obtained the number of publications for each gene studied here from PubMed and compared it to each gene length group and with the number of interactions (Fig 7B). We observed that the number of interactions and publications were significantly different between each gene length group (Wilcoxon rank sum test, p-value < 2.2E-16 for both comparisons), with both being higher for the group comprising of longer length genes. In order to assess the level of influence of publication bias in our protein-protein interaction dataset, we used correlations between the values of protein-protein interactions and the number of publications and we observed that, for both gene length groups, the correlations were not the strongest (Kendall test; Longest genes, tau = 0.26, p-value < 2.2E-16; Smallest genes, tau = 0.36, p-value < 2.2E-16), implying that while there might be some publication bias in effect, the strength of that effect is rather weak.

However, for the group of the longest genes, 208 (21%) entries were of zero value, while for the smallest group of genes, 544 (55%) entries were of zero value. This means that there were either no physical interactions for those genes, or that there were no entries in BioGRID for them. In order to account for this, and similarly to what we did for the co-expression analysis, we extracted the top 100 genes with the most and fewest protein-protein interactors (without null values) in our dataset and we observed the distribution of their transcript length. We observed that genes with the highest protein-protein interactions were longer (median transcript length = 3737), than genes with the lowest amount of protein-protein interactions (S10B Fig, Wilcoxon rank sum test, p-value = 0.039, median transcript length = 2764).

## Discussion

With this work, we tried to elucidate what factors affected gene length and whether gene length had a role in determining the function of their proteins in the cell. Even looking at the 5 longest genes, we can get a small glimpse into one these objectives. *TTN* is the longest transcript in the human genome, and serves several important functions in the skeletal and cardiac muscles, and is often involved in structure, sensory and signalling responses [20,66,67]. The mucin *MUC16* (or CA125) is mostly known as a biomarker in ovarian cancer and is used to monitor patients as an indicator of cancer recurrence [21,68,69]. SMAD family member 2 (*SMAD2*) is thought to play a critical role in neuronal function [17] and to have a protective role in hepatic fibrosis [70]. The gene *FLRT2* is believed to have a role in tumour suppression in breast and prostate cancer [71, 72] and, in mice models, *FLRT2* has been found as a guiding agent in neuronal and vascular cells [18, 73]. For the *GRIN2B* gene, it has been shown to play an important role in the neuronal development and cell differentiation in the brain [19, 74]. We cannot obtain any information at the moment pertaining to the function of the 5 smallest genes, since all of them are either novel and have yet to be properly studied, or could be annotation errors in the assembly.

In order to deeply understand the effects of gene length in protein function, we performed a functional analysis. For longer length genes, the GO terms obtained were mostly associated with neurons, for example terms like axon development, axon part, neuron to neuron synapse, actin and cell polarity [75] and GTPases [75]. For tissue specific genes, brain and nerve had the longest genes. Looking at the KEGG Pathways associated with the longest genes, the categories present are in the brain, cancer, heart diseases and muscle. Previous studies have associated longer length genes with neurons [76, 77] and muscle [78]. Due to the very nature of longer genes, one expects high rates of mutation, not only due to their size, but also due to possible collisions between the RNA polymerase and the DNA polymerase, which causes instability and possible mutations [79]. It is not surprising to find associations between longer genes with cancer [15] and hearth pathologies often caused by mutations in particularly long genes, like *DSC2* and *TTN* [80–82].

Looking at our smaller genes group, most of the GO terms provided were associated with the skin, for example skin development and cornified envelope, or with the immune system, for example, defence response to other organism and receptor agonist activity. Smaller tissue specific genes also have a major presence in the skin. With regards to the KEGG Pathways associated with the smaller genes, most pathways were involved in the immune system, with a few also being present in neurodegenerative diseases and in the skin. Previous studies have observed that most genes associated with immune functions are rather small in size [83]. However, there are no studies to support the association of smaller genes with skin development. The categorization on the basis of published work has its advantages, but there is often overlapping of functions within these categories, for example, calcium signalling also happens in the muscle [84] and immune system [85], Wnt signalling pathway also has a role in cancer [86], TGF-beta signalling pathway can also be associated with the immune system [87], among others. In spite of this, our findings lead us to believe there is a disparity in gene sizes for genes that have a role or are present in tissues with very little to almost no development pos-natally (like neuron) and genes (not involved in housekeeping) that are quite frequently expressed during a human’s whole lifetime (like in skin development and immune response) or involved in providing functions with fast responses. Corroborating with our findings for the functional analysis, a recent preprint has showed that, with age, there is a downregulation of long transcripts and an upregulation of short transcripts, in a phenomena they named “length-driven transcriptome imbalance”, which in humans it affects the brain the most [88]. As we observed, smaller genes can be associated with the immune system and inflammation has a role in many ageing-related diseases [89], while longer genes are mostly associated with brain development, a function that happens early in life.

To understand whether there were factors that had an influence in gene length, we performed several correlation analysis. Overall there was no really strong correlation observed between the gene characteristics studied and transcript length. The biggest significant positive correlations were with protein size and number of SNPs, with transcript count, number of introns, GC content, and average gene expression having a weak significant positive correlation. Results of the correlation between average gene expression and transcript length were not in line with previous observations, which suggested that highly expressed genes are often smaller in length [11]. We also observed that among smaller genes, the average gene expression was, in fact, the highest (S4D Fig). However, genes with smaller lengths also had a great variability in the average gene expression values, and there was almost no correlation between transcript length and average gene expression. What has been stated in the previous studies is relevant, but the whole image is not captured properly. Rather than stating that the smaller genes are highly expressed, it is more accurate to say that smaller genes have a greater variability of levels of expression than longer genes. Similar to the correlation results for number of SNPs, both synonymous and missense mutations were also highly correlated with transcript length. It is particularly interesting that the correlation values were so high for missense mutations, since these may cause loss of function in the resulting protein. Likewise, it could be one of the reasons why the correlation between nonsense mutations and transcript length is weaker than the other two. Other works [9] have used the MIS/SYN and NONS/SYN ratios as a measure of functional importance, and we can, albeit faintly, observe here that longer genes appear to be more functionally important than smaller gene. The negative correlation between these ratios showed that longer genes may have more mechanisms in place to prevent loss of function mutations, when compared with synonymous mutations. Moreover, we also have to take account of “outliers” when looking into the correlation between transcript length and protein size (S4C Fig), specifically for longer genes. One would expect that for longer genes, the proteins produced would have a size comparable to their length and not be extremely small. However, after observing these outliers and we found that their protein size was rather small due to the presence of very long 3’UTR regions. While these regions still account for the calculation of gene size, they are not translated into the protein, causing the presence of these “outliers”. Previous studies have shown that the brain has a preference for these long 3’UTR regions [90, 91].

Interestingly, we also noticed that genes associated with ageing tend to be longer than the rest of the protein-coding genome. Moreover, we also showed that the overall (not tissue dependent) expression of genes with age appears to disregard transcript length, and that the brain seems to favour the expression of smaller genes with age. This last result, seems on par with the previously mentioned observations by Stoeger et al. [88], where they also witnessed the upregulation of smaller transcripts with age, especially in the brain. However, the results pertaining to the overall expression of genes with age seems to be different between what Stoeger et al. observed, with transcript length as an important source of ageing-dependent changes in values of expression, and what we observed based on Palmer et al. signatures of ageing [60], where transcript length does not influence the expression of genes with age. It is possible that these two works found two different sets of genes whose expression is affected in the ageing process. As such, further works should prove useful in dictating whether or not transcript length plays a major role in the expression of genes with age.

When comparing gene length with the dN/dS ratio for three organisms (Gorilla, Chimpanzee and Mouse), longer genes appeared to evolve under constraint, while for smaller genes there was a promotion for changes in the genes by natural selection. Previous studies have shown that, for genes classified as “old” (by virtue of having orthologues in older organisms), their length will be longer, they will have more introns and they evolve more slowly than smaller genes [7, 8]. In terms of the co-expression analysis and protein-protein interactions, the longer genes, in general, had the most co-expression partners and protein-protein interactions. Further validating our observations, we also saw that top hundred highest co-expression genes and PPI were longer in length as compared to lowest co-expression genes and PPI.

As a result of this work we have noticed that not all genes are studied with the same depth. Some genes have more information related to expression or function than others. We observed this especially within our 5% list of longest and smallest genes. Longer length genes had more functional information readily available than smaller ones. We can also observe that in the publication bias analysis for protein-protein interactions, where genes with longer lengths had more publications than smaller genes. Indeed, other groups have found that gene length can be an important predictor of the number of publications, and that novel genes are not often studied to their full capacity [92], while others have found that genetic associations tend to be more biased towards longer genes [93, 94].

The present study has its own limitations. One of the limitations for this sort of study is that, the results might be “time-specific”. With new discoveries related to the human genome and its genes, the trends here observed might change, specifically when it concerns the currently extremely untapped field of smaller genes. Similarly as we previously noted, longer genes have a lot more information related to them, when compared with their smaller counterparts. While our findings with respect to the longer genes might be mostly reliable, we cannot show the same confidence in case of the smaller genes, considering that a lot of these genes were novel and have yet to be properly studied. However even after taking account of the above limitations, the present study still provides some very interesting insights pertaining to gene length and its possible role in early life development, diseases and response time in the human genome.

## Conclusion

With this work we aimed to better understand the effects of gene length in gene function and factors that affected it. We observed that, for most of the factors studied, there was not a particularly strong correlation with transcript length. The strongest correlations here detected were associated with the number of SNPs and the protein size. We also showed that, for smaller genes, its association with high levels of expression is not entirely correct and that, instead, there is great variability of expression values among them. We also observed that longer genes appear to have the most co-expression partners and protein-protein interactions, in comparison to their smaller counterparts.

In case of the functional analysis, we observed that longer genes favoured functions in the brain, cancer, heart and muscle, while smaller genes are strongly associated with the immune system, skin and neurodegenerative diseases. This lead us to believe that gene length could be associated with the frequency of usage of the gene, with longer genes being less often used past the initial development and smaller genes playing a frequent role daily in the human body.

## Methods

### Data retrieval and filtering

All protein-coding human transcripts and genes (N_transcripts_ = 92696), their length, transcript count and GC content were obtained using the biomart [16] website (GRCh38.p12, Ensembl 96, April 2019). Transcript length is defined by Ensembl as the total length of the exons in a gene plus its UTR regions lengths. Gene length was obtained using the R (version 3.5.2) package EDASeq (version 2.14.1). Using R, the transcripts with the highest transcript length per gene were selected. In case of ties, due to multiple transcript having the same length per gene, we used some tags (APPRIS annotation was the principal one, if there was an entry in RefSeq or GENCODE) used by ensemble as a tie-breaker. Should that fail, the oldest transcript was chosen, by means of having a smaller numerical ID. Transcripts associated with PATCH locations or assemblies were removed from our dataset. For each transcript, we obtained data regarding their number of exons, CDS length, number of SNPs, synonymous (“synonymous_variant”), missense (“missense_variant“) and nonsense (“stop_gained”) SNPs, protein length, dN and dS values, using the biomart (version 2.38.0) package in R. For the dN and dS values, only values associated with One to One orthologues were selected for the present analysis. Average expression was obtained from the USCS Table browser tool [95], using expression as the group and the GTEx Gene track. Tissue specific Tau values of expression were obtained from a previous work [60]. The number of SNPs per gene was obtained using the Ensembl API, R and the httr (version 1.4.0) and jsonlite (version 1.6) packages.

The whole file produced and used in the analysis for this work can be found on the Supplementary Table 1 (N = 19714).

Gene names of genes related with ageing (N = 307) were obtained from GenAge (Build 19) [62].

### Statistical tests, graphs and other packages

R and the function corr.test were used to perform the correlation tests. Due to the abundance of the data, there were a lot of ties in the ranks, which prevented the usage of Spearman’s correlation, so instead we chose to use the Kendall test for the correlations. The figures produced in this work were created using the ggplot2 (version 3.2.0) package in R. Other packages used over the course of this work were: corrplot (version 0.84), psych (version 1.8.12), ggpubr (version 0.2.1), stringr (version 1.4.0), dplyr (version 8.0.1), plyr (version 1.8.4) and tidyr (version 0.8.3).

### Functional Analysis

WebGestalt (2019 release) [22] was used to do the Overrepresentation Enrichment Analysis for each of the gene ontology categories (Biological Process. Cellular Component and Molecular Function). The top 5% genes, with the highest and lowest gene length, were ran against the reference option of genome. The significance level was FDR<0.05 and the multiple test adjustment was done using the Benjamini–Hochberg method.

For confirmation of the results, the same two 5% lists were run on DAVID’s [23, 24] annotation clustering option, using the complete human genome as background. Only terms with p-value and FDR smaller or equal to 0.05 were considered. Default categories were used except for the category “UP_SEQ_FEATURE”, since it was introducing a lot of redundant results.

To help better visualize the GO terms obtained from the analysis above described, the tool REViGO [59] was used. The p-values here considered were the FDR values obtained previously, with the human database option used for the GO terms.

In regards to the analysis done using the KEGG pathways, the grouping of genes and pathways was obtained from the Molecular Signature Database (version 6.2) [26,27,96–99], like it was done previously by another group [15]. Additionally, the colouring of the box plot was done based on the fact that the pathway in question is directly associated with the category (when the KEGG Pathway schematic shows cells from the category) or if they could be indirectly associated with the category (using available literature). For this last case, appropriate literature was selected if they mentioned elements of the KEGG Pathway being involved in said category.

### Co-Expression Analysis

Co-expression correlation values were extracted from GeneFriends [64]. For each gene (N = 19714), in the whole dataset and in the top 5% lists of genes with the longest and smallest transcript length (N = 986 for each list), the number of genes with correlation values superior or equal to 0.6 or smaller or equal to -0.6 were obtained using R. From our original dataset (N=19714 genes), 1046 genes were not present in GeneFriends (whole dataset), of which, 25 missing genes were within the High group and 110 missing genes were within the Low group. For obtaining the median values of genes present in the GeneFriends database, the co-expression values for each gene across the database were merged and this was followed by calculation of median values using R.

### Protein-Protein Interaction Analysis

BioGRID (release 3.5.174) REST API [65] in conjugation with the R package httr was used to obtain all protein-protein interactions for the whole dataset and for the top 5% longest and smallest genes. All redundant and genetic interactions were removed from this analysis.

For the publication bias, the number of publications, in PubMed, per gene of each group was obtained using the Entrez Programming Utilities (E-utilities), and the R packages XML (version 3.98-1.19), httr and biomart.

## Acknowledgements

The authors wish to thank past and present members of the Integrative Genomics of Ageing Group for useful suggestions and discussion, in particular Kasit Chatsirisupachai and Daniel Palmer.

## Supporting information

**S1 Table. Dataset with the highest protein-coding transcript length per Gene, in human.**

**S2 Table. Functional analysis results for WebGestalt and DAVID.**

**S3 Table. KEGG Pathway IDs used in Supplementary Figure 2.**

**S4 Table. Co-Expression results.**

**S5 Table. Number of Protein-Protein interactions and Publications in Pubmed for each gene in the dataset.**

**S1 Fig. Functional analysis results for Cellular Component and Molecular Function.**

**S2 Fig. Transcript length distribution per KEGG Pathway.**

**S3 Fig. Correlation results for Number of SNPs, protein size, transcript count, GC content and synonymous, missense and nonsense mutations against transcript length.**

**S4 Fig. Gene length and intron distribution in the human genome.**

**S5 Fig. Transcript length distribution for genes specifically expressed in the given tissues.**

**S6 Fig. Transcript length distribution for ageing related genes and for the rest of the dataset.**

**S7 Fig. Evolution results for mouse, gorilla and chimpanzee.**

**S8 Fig. Co-expression results.**

**S9 Fig. Protein-protein interactions results.**

