## Supplementary material for "Gene size matters: What determines gene length in the human genome?": S1 Fig

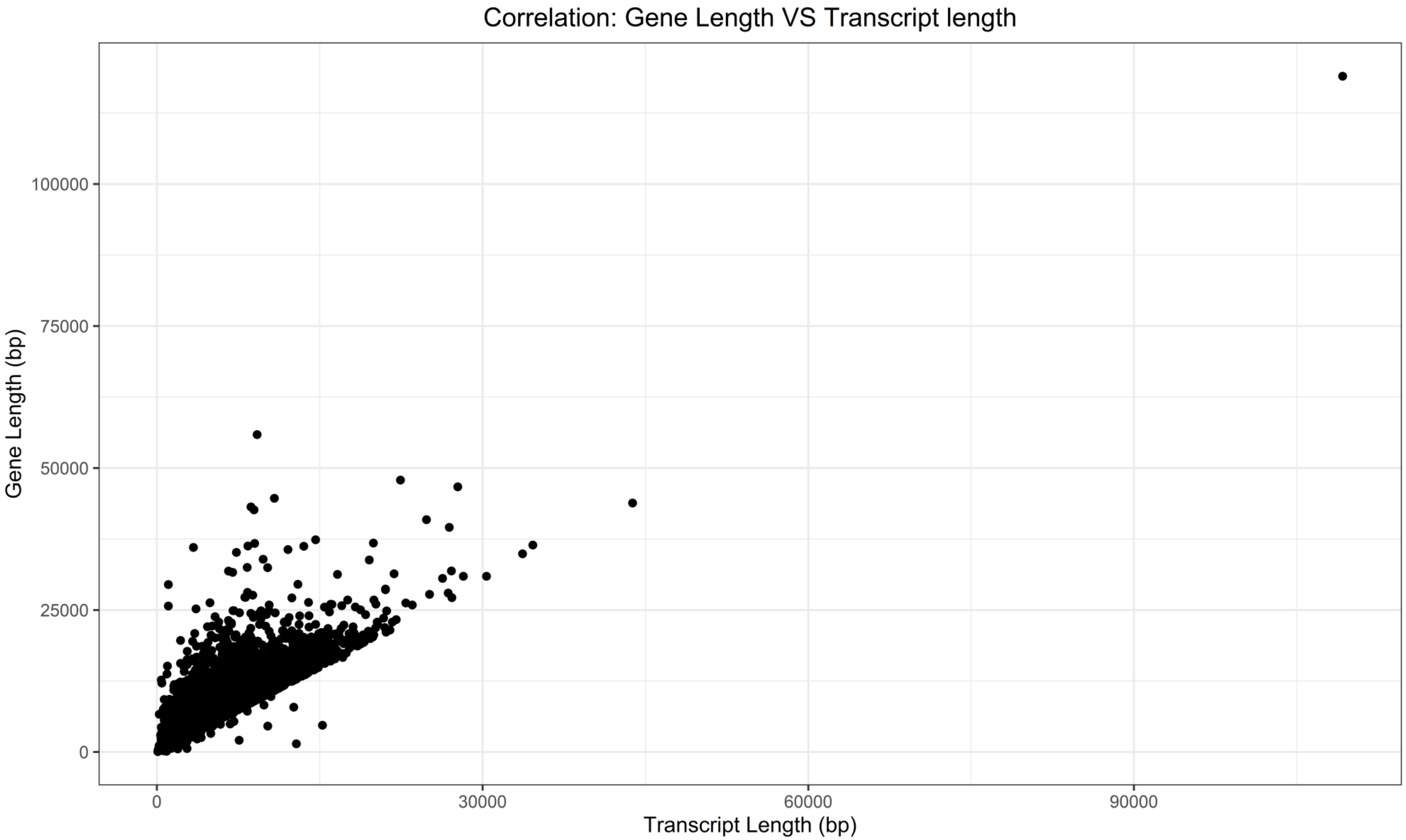


**Supplementary Figure 1.**
Correlation between Gene Length and Transcript Length from the longest transcript in a gene. Gene Length was obtained using the EDASeq package and the Transcript Length was obtained using biomart.
