## Supplementary material for "Gene size matters: What determines gene length in the human genome?": S2 Fig

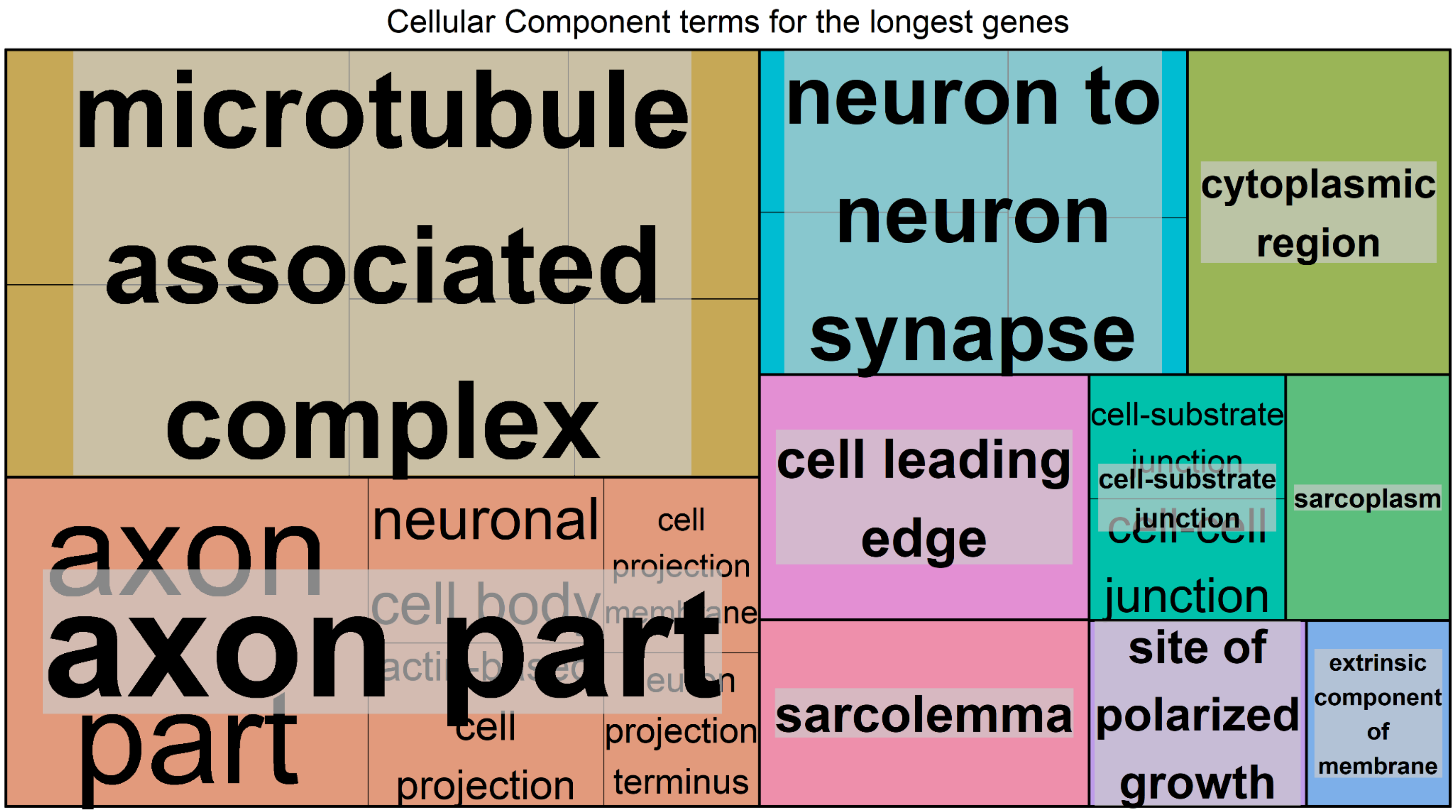


**Supplementary Figure 2A.**

Cellular Component terms found associated to genes with the longest transcript length. The significance level was $p<0.05$ and the FDR was set at 0.05. FDR estimation was done using the Benjamini–Hochberg method.

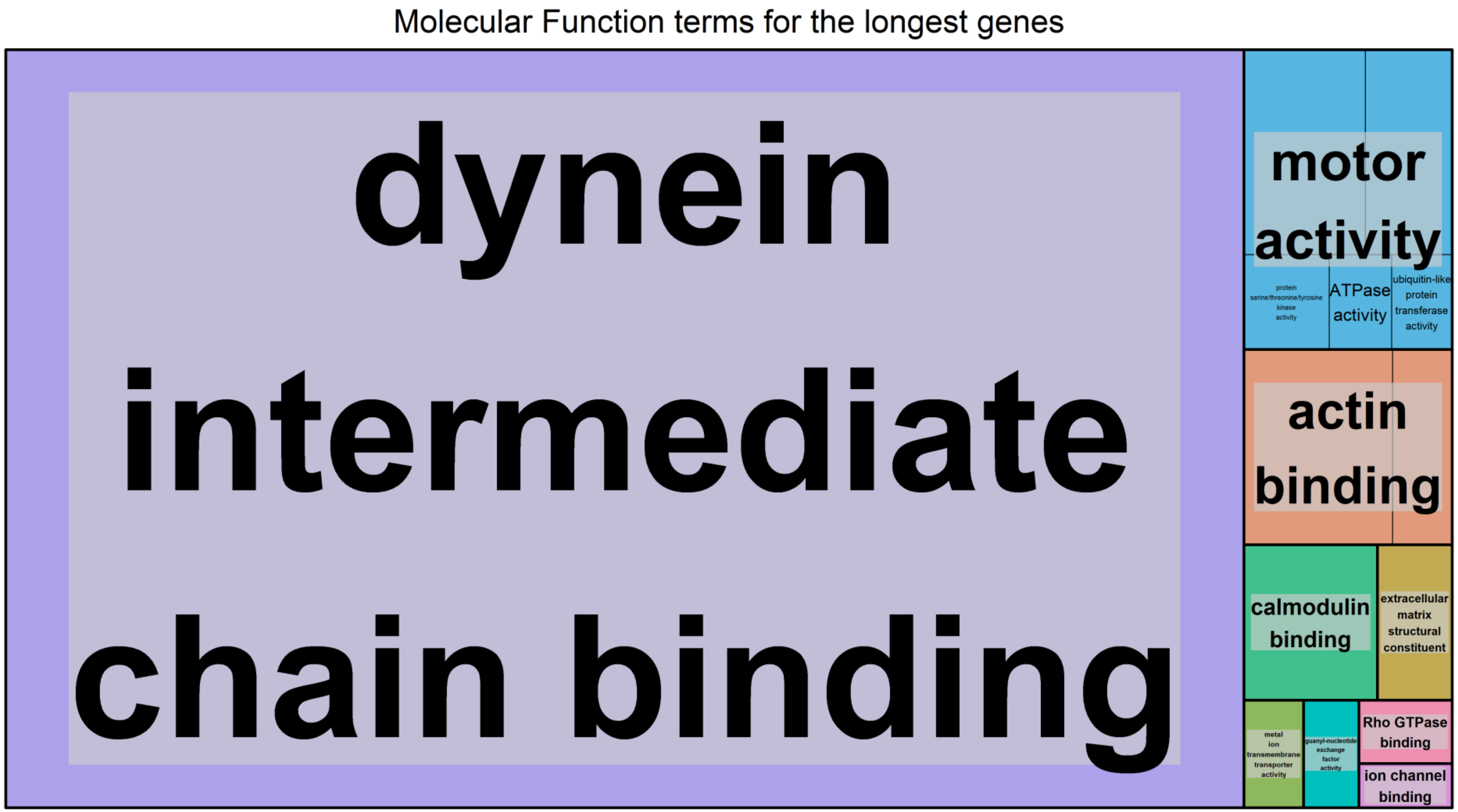


**Supplementary Figure 2B.**
Molecular Function terms found associated to genes with the longest transcript length. The significance level was $p<0.05$ and the FDR was set at 0.05. FDR estimation was done using the Benjamini–Hochberg method.

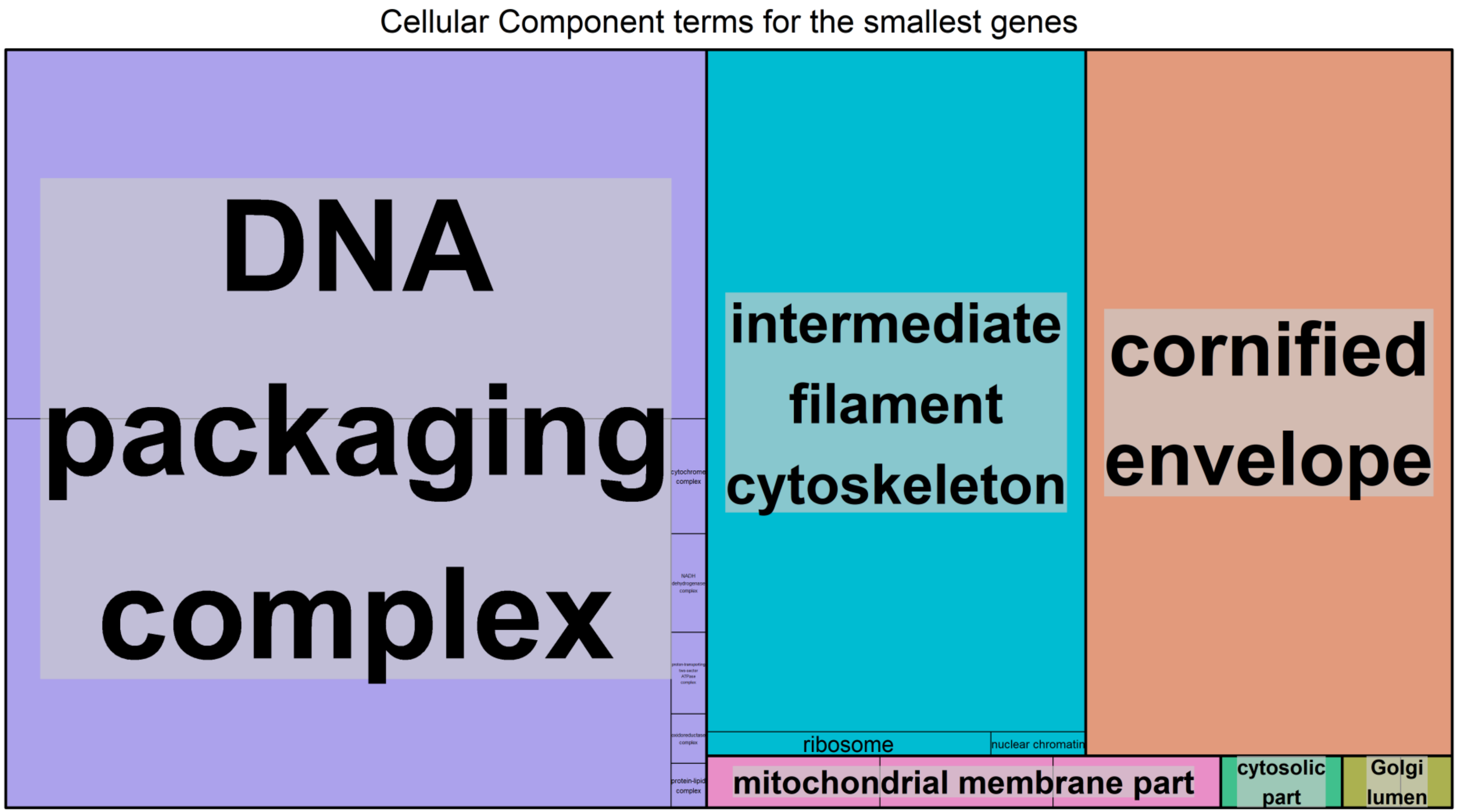


**Supplementary Figure 2C.**
Cellular Component terms found associated to genes with the smallest transcript length. The significance level was $p<0.05$ and the FDR was set at 0.05. FDR estimation was done using the Benjamini–Hochberg method.

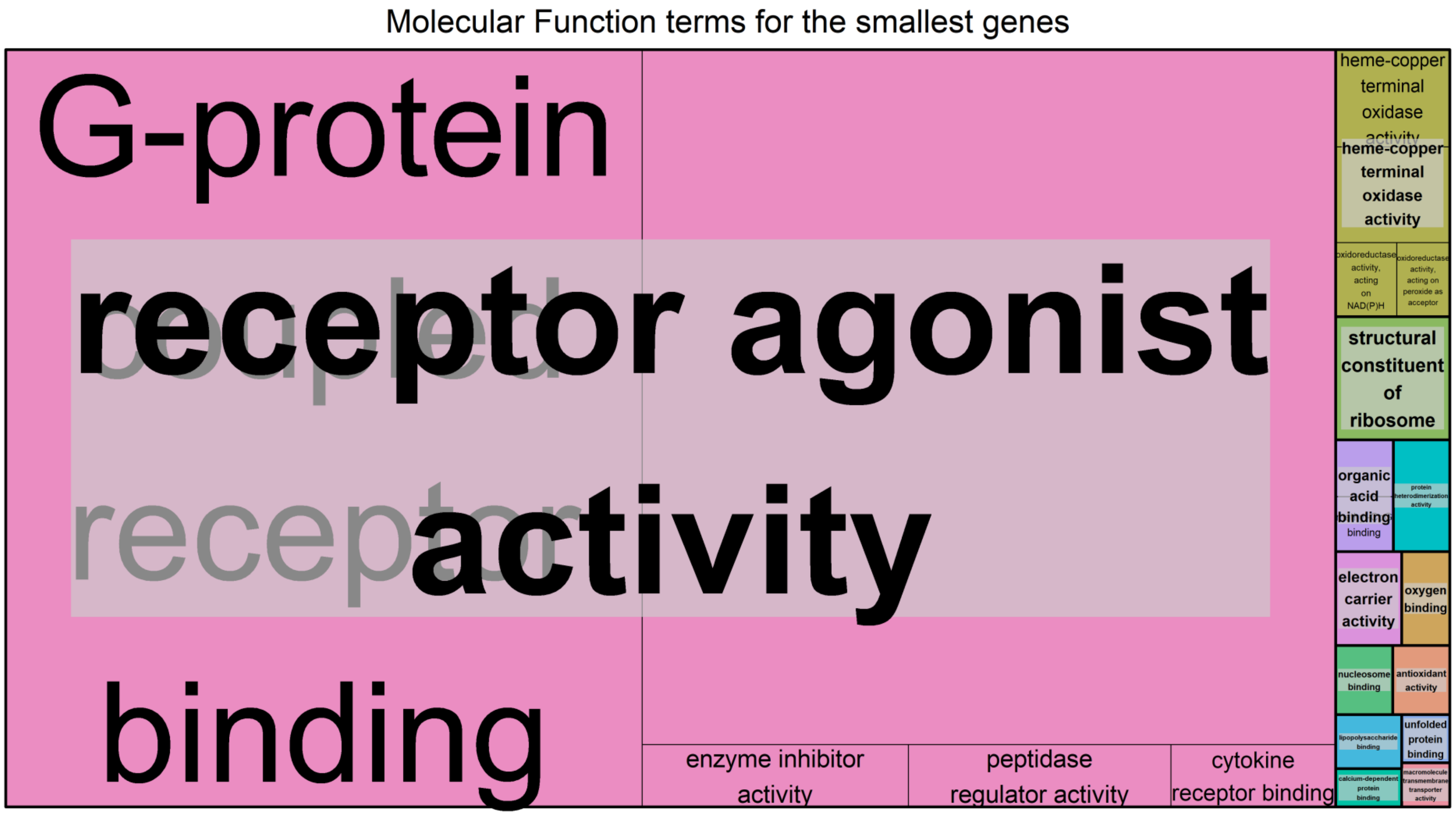


**Supplementary Figure 2D.**
Molecular Function terms found associated to genes with the smallest transcript length. The significance level was $p<0.05$ and the FDR was set at 0.05. FDR estimation was done using the Benjamini–Hochberg method.
