## Supplementary material for "Gene size matters: What determines gene length in the human genome?": S3 Fig

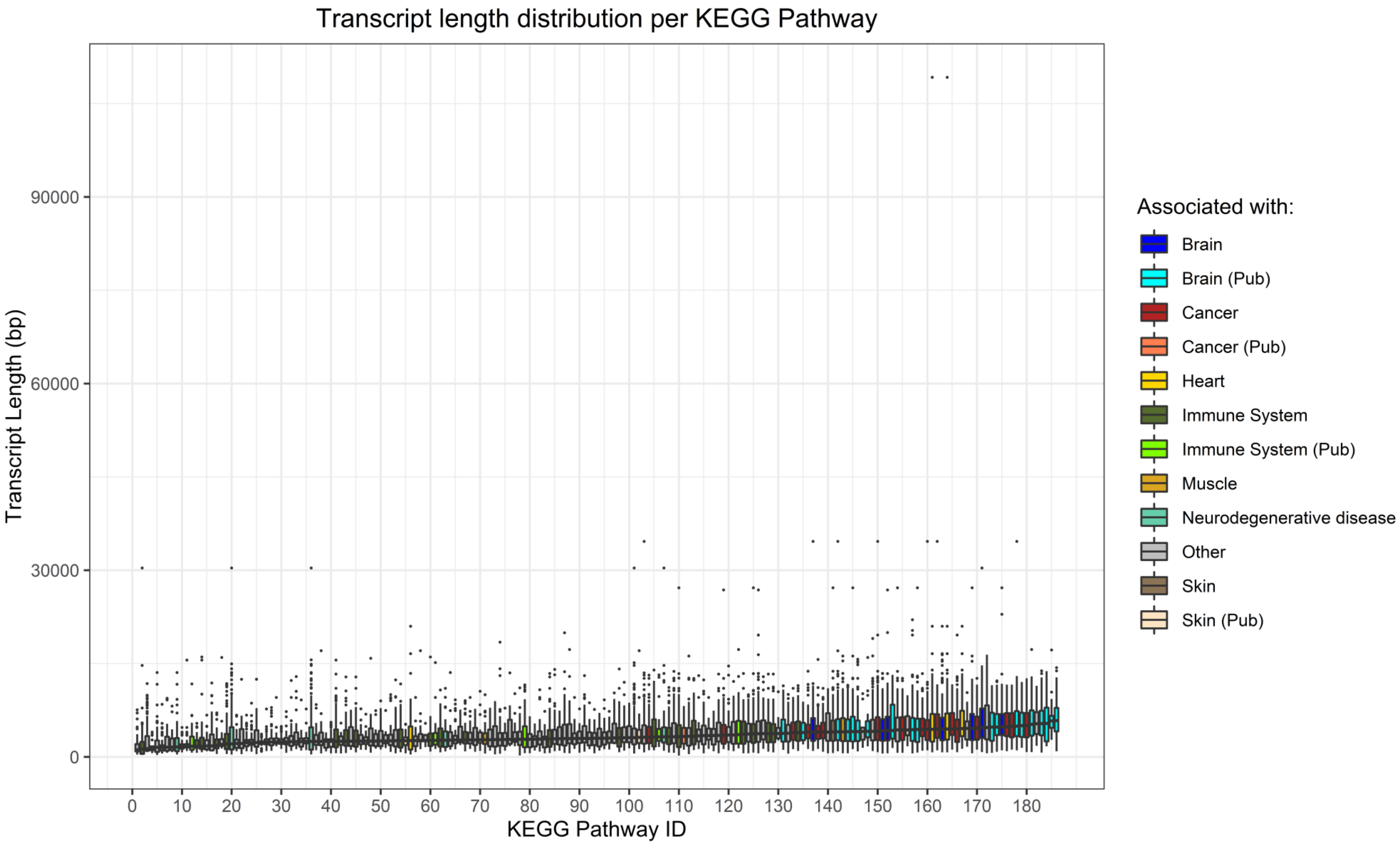


**Supplementary Figure 3A.**
Transcript length distribution per KEGG Pathway. Colours illustrate what the KEGG pathway has been directly associated with, due to it being stated in the pathway itself, or indirectly associated with (Pub tag), by means of literature references. KEGG Pathway IDs can be found in the Supplementary Table 3. KEGG Pathways and genes involved in said pathways were obtained from the Molecular Signature Database.

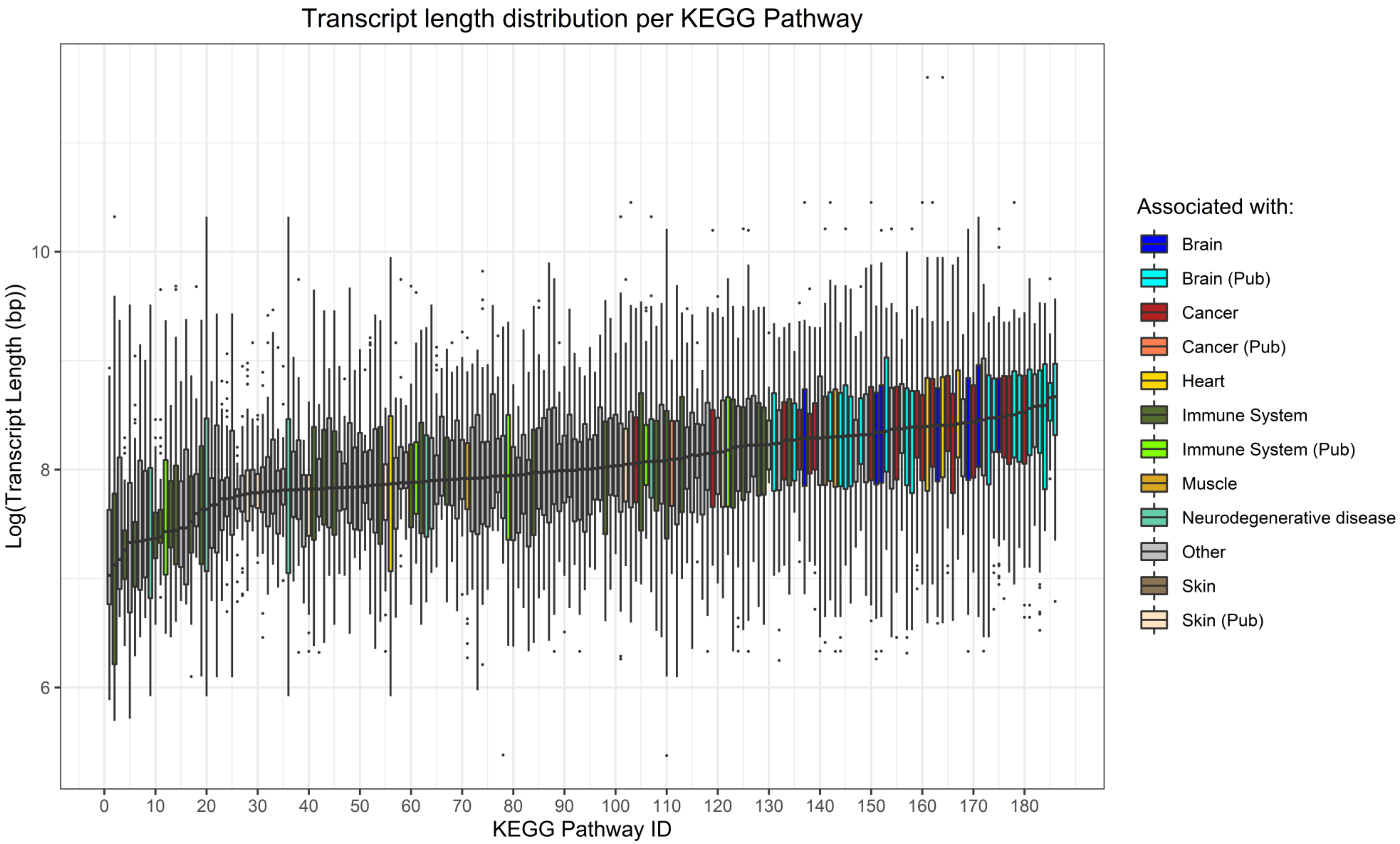


**Supplementary Figure 3B.**

Log transformed transcript length distribution per KEGG Pathway. Colours illustrate what the KEGG pathway has been directly associated with, due to it being stated in the pathway itself, or indirectly associated with (Pub tag), by means of literature references. KEGG Pathway IDs can be found in the Supplementary Table 3. KEGG Pathways and genes involved in said pathways were obtained from the Molecular Signature Database.
