## Supplementary material for "Gene size matters: What determines gene length in the human genome?": S4 Fig

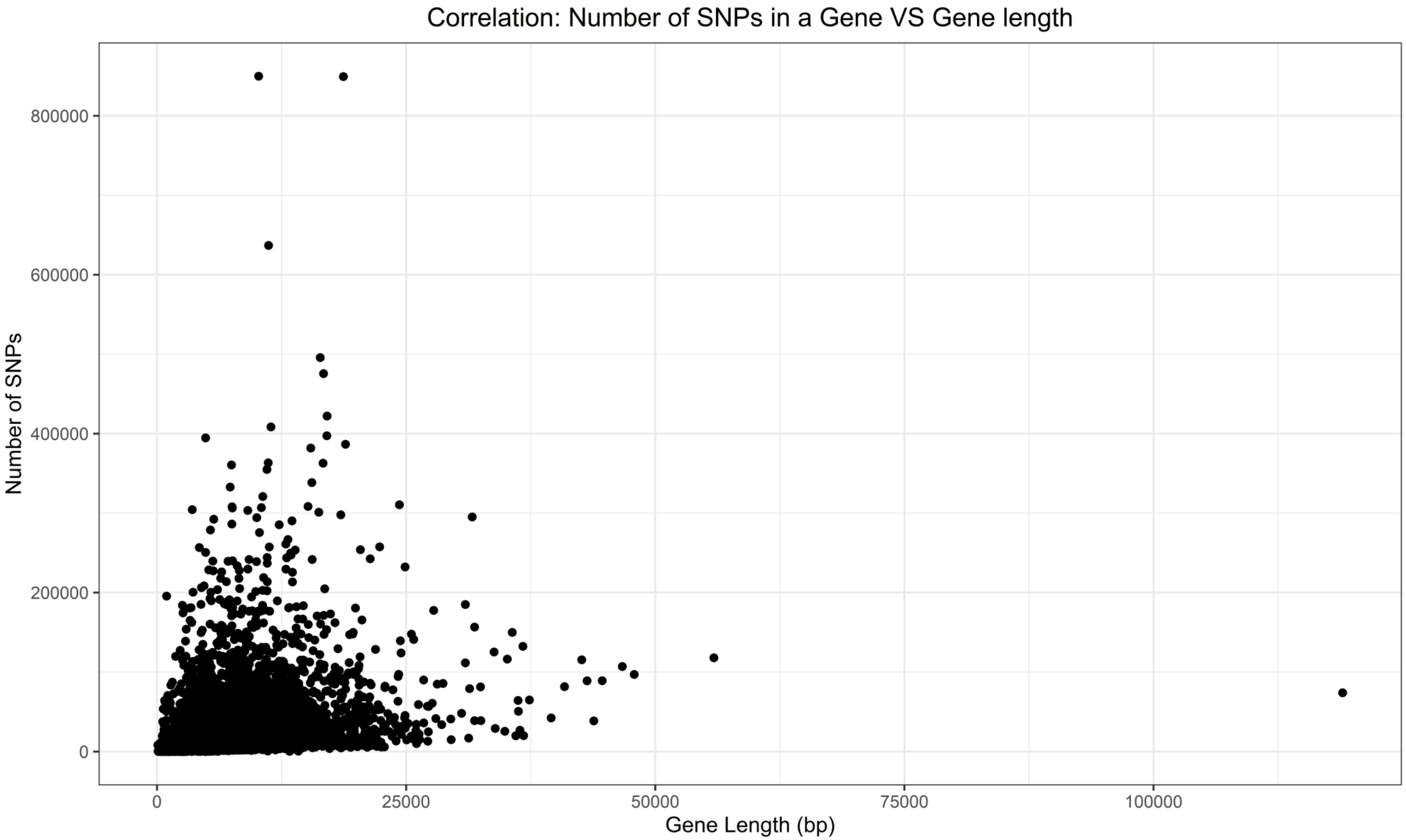


**Supplementary Figure 4A.**
Correlation between the Number of SNPs per gene and Gene Length. Gene Length was obtained using the EDASeq package and the Number of SNPs was obtained using the Ensembl API.

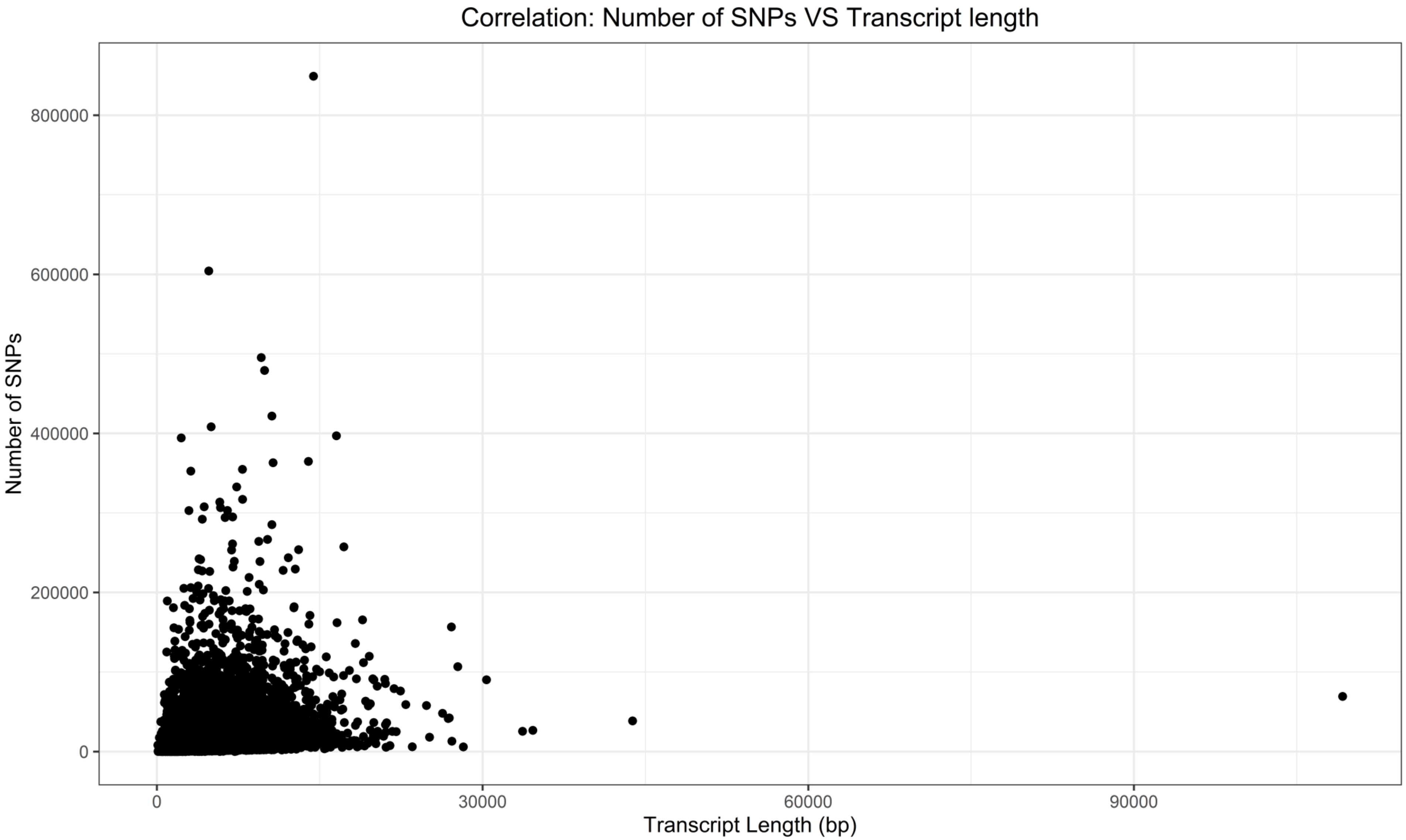


**Supplementary Figure 4B.**
Correlation between the Number of SNPs and Transcript Length. Number of SNPs and Transcript Length were obtained using biomart.

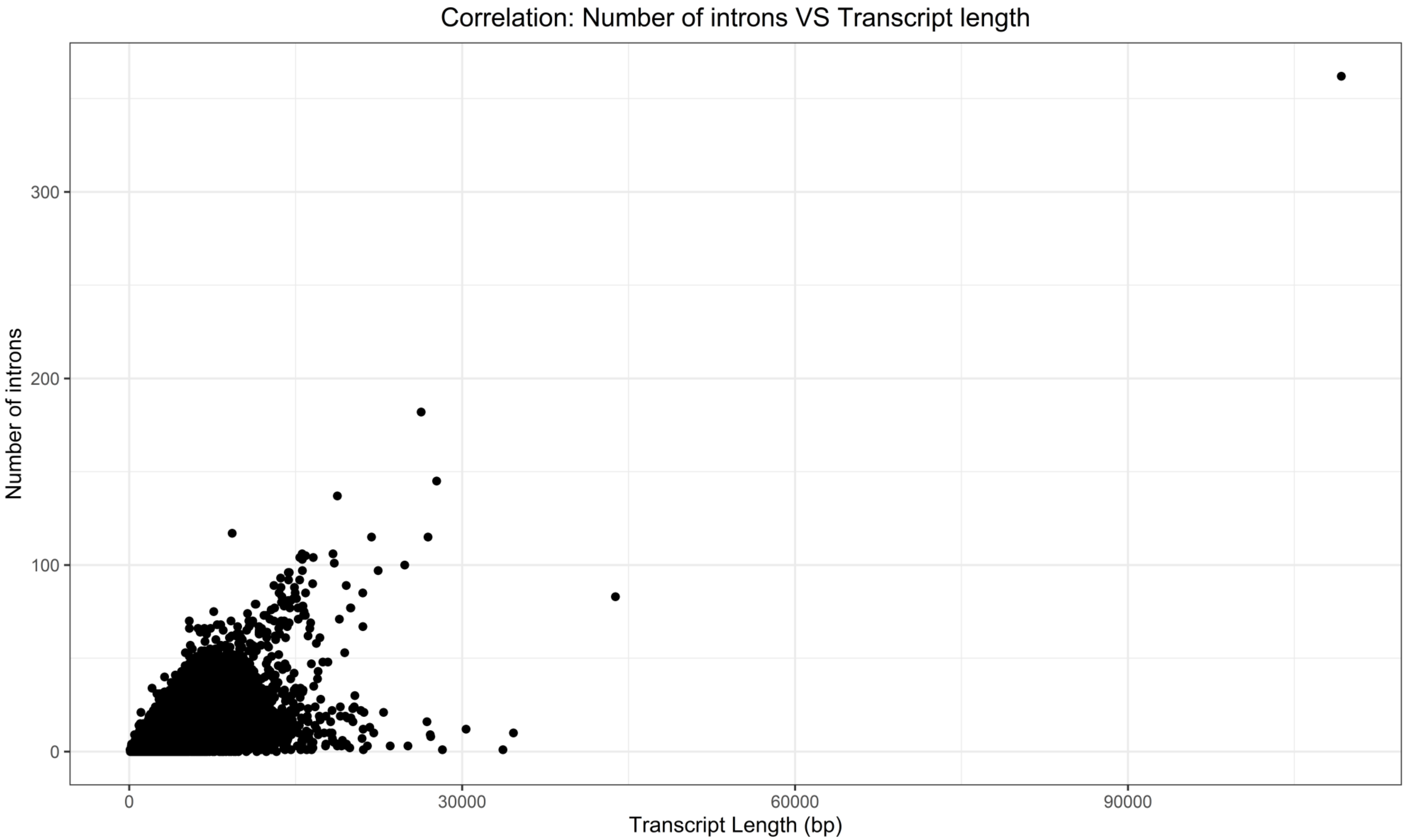


**Supplementary Figure 4C.**
Correlation between the number of introns and Transcript Length. Number of introns and Transcript Length were obtained using biomart.

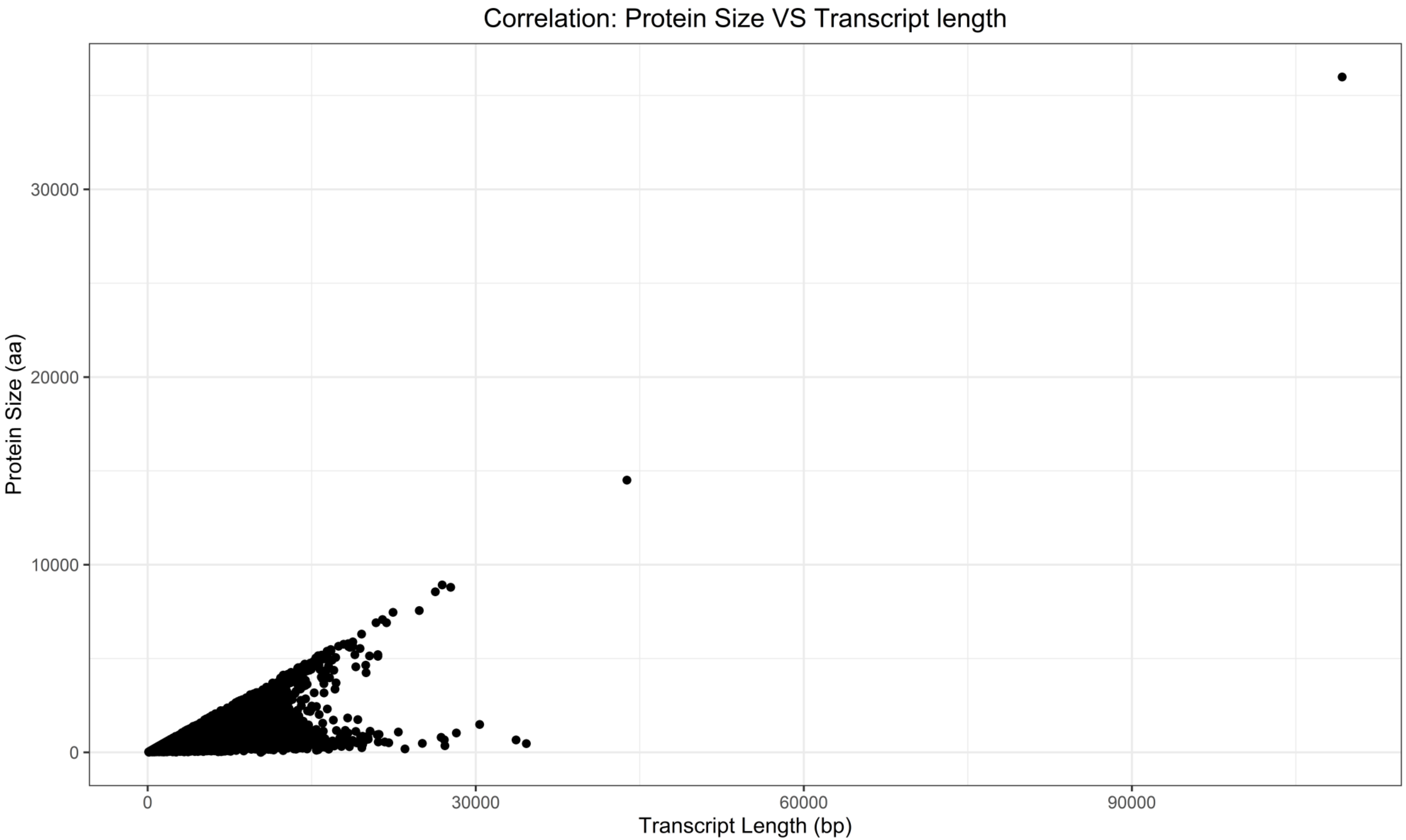


**Supplementary Figure 4D.**
Correlation between Protein size and Transcript Length. Protein size and Transcript Length were obtained using biomart.

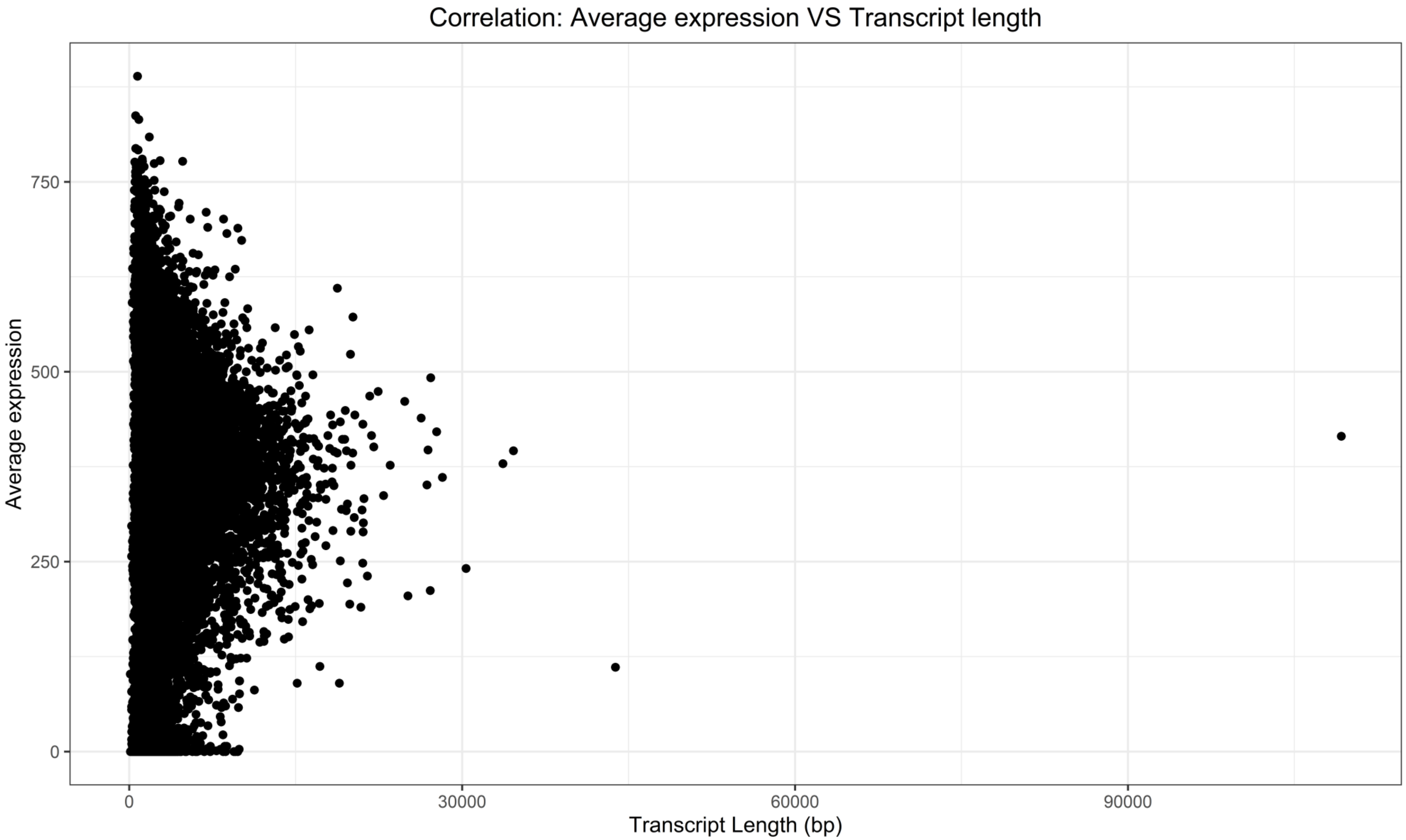


**Supplementary Figure 4E.**
Correlation between the Average Gene Expression and Transcript Length. Average Gene Expression was obtained from the UCSC Genome browser and Transcript Length was obtained using biomart.

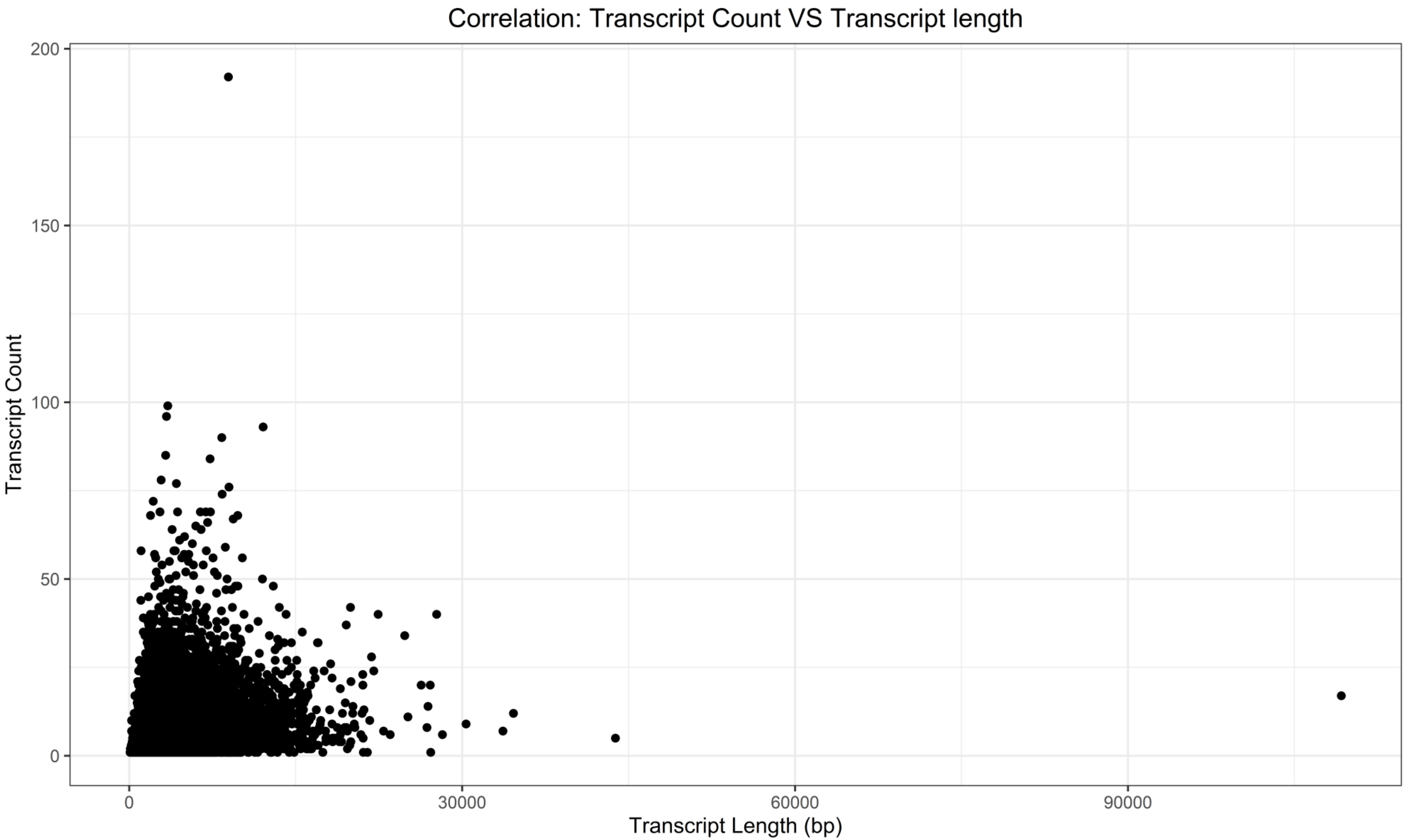


**Supplementary Figure 4F.**
Correlation between Transcript count and Transcript Length (bp) (Kendall test, tau = 0.22, p-value < 2.20E-16). Transcript count and Transcript Length for each transcript were obtained using biomart.

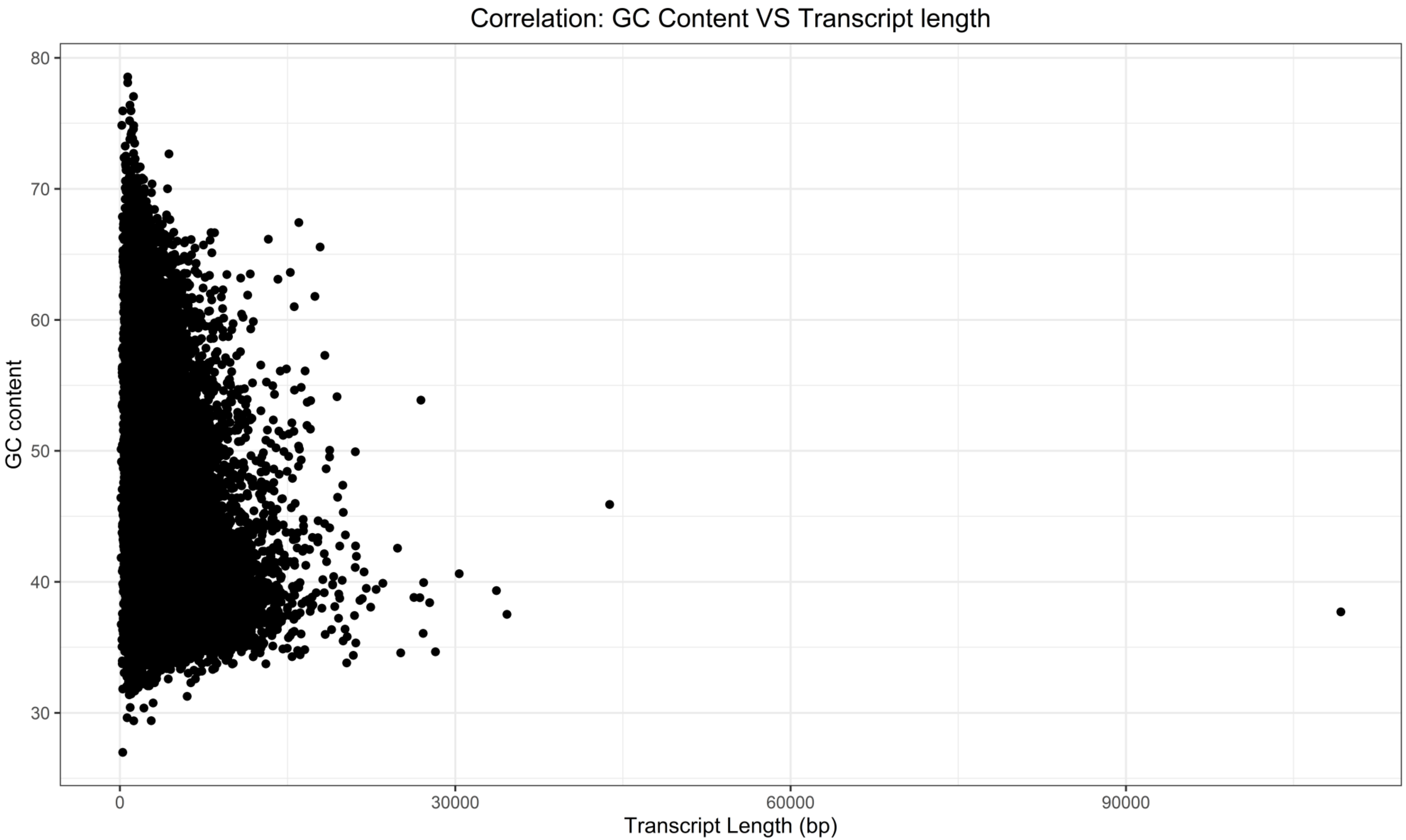


**Supplementary Figure 4G.**

Correlation between the GC Content and Transcript Length (bp) (Kendall test, tau = -0.19, p-value < 2.20E-16). GC Content and Transcript Length were obtained using biomart.

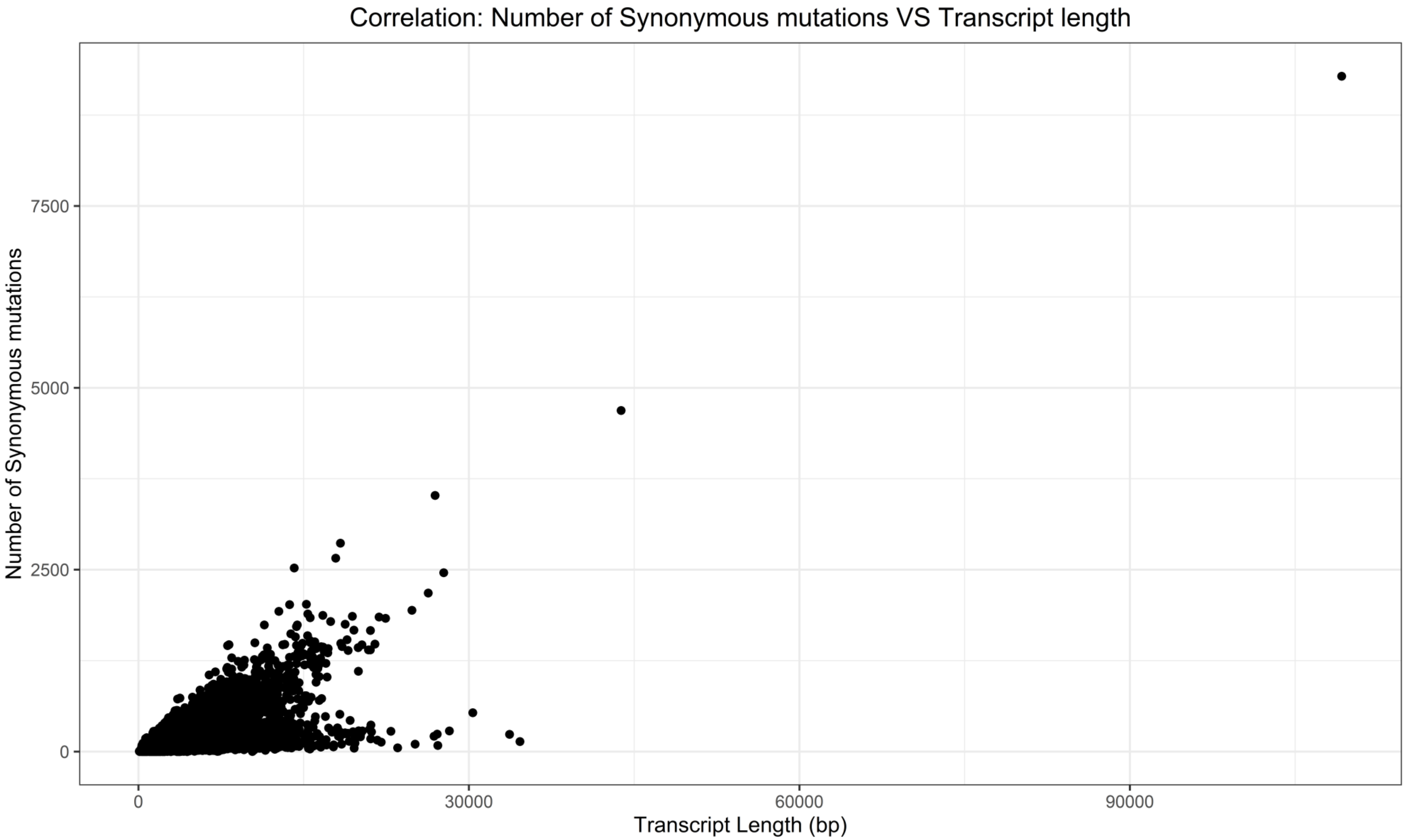


**Supplementary Figure 4H.**
Correlation between synonymous mutations and Transcript Length (bp) (Kendall test, tau = 0.44, p-value < 2.20E-16). The number of synonymous mutations and Transcript Length were obtained using biomart.

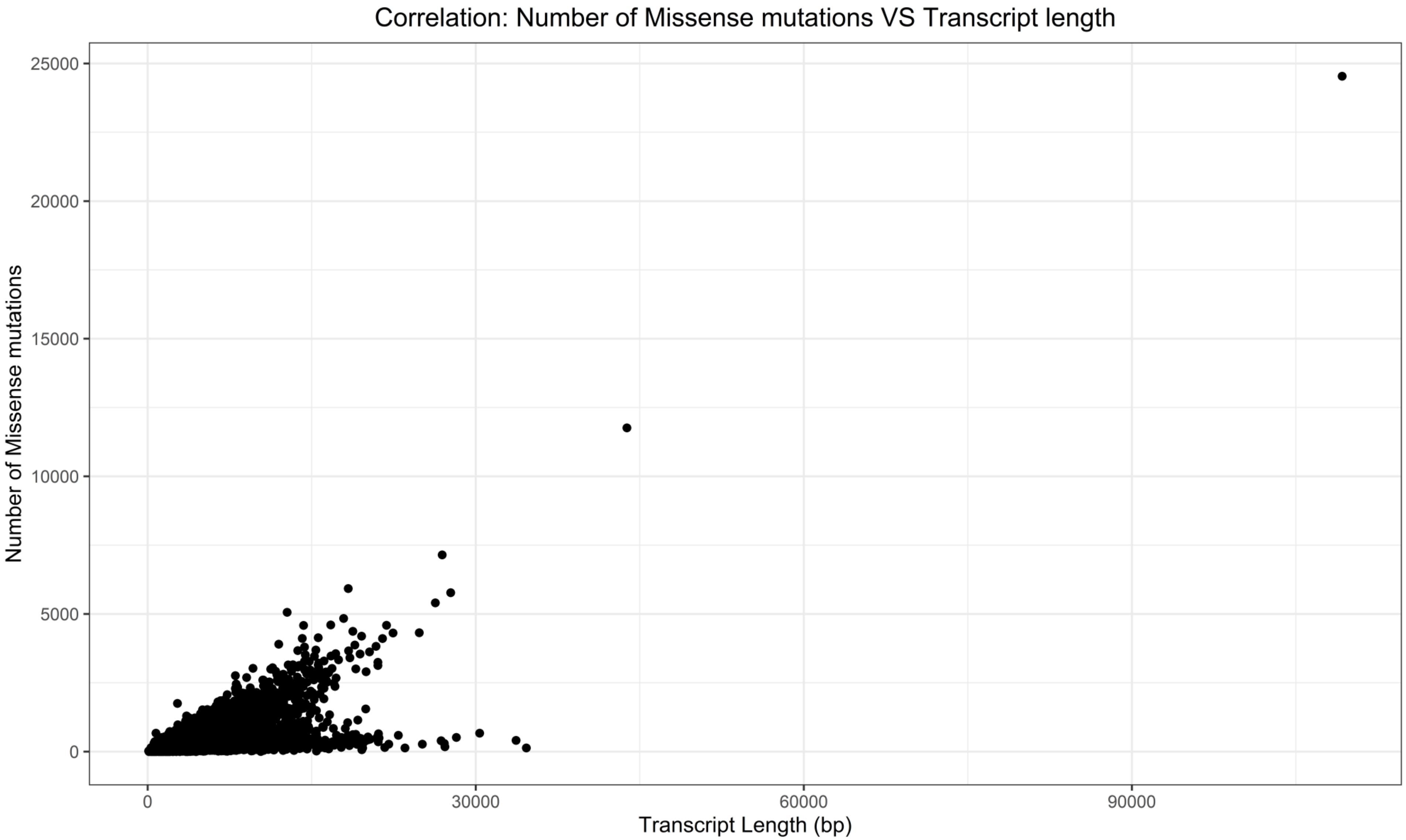


**Supplementary Figure 4I.**
Correlation between missense mutations and Transcript Length (bp) (Kendall test, tau = 0.42, p-value < 2.20E-16). The number of missense mutations and Transcript Length were obtained using biomart.

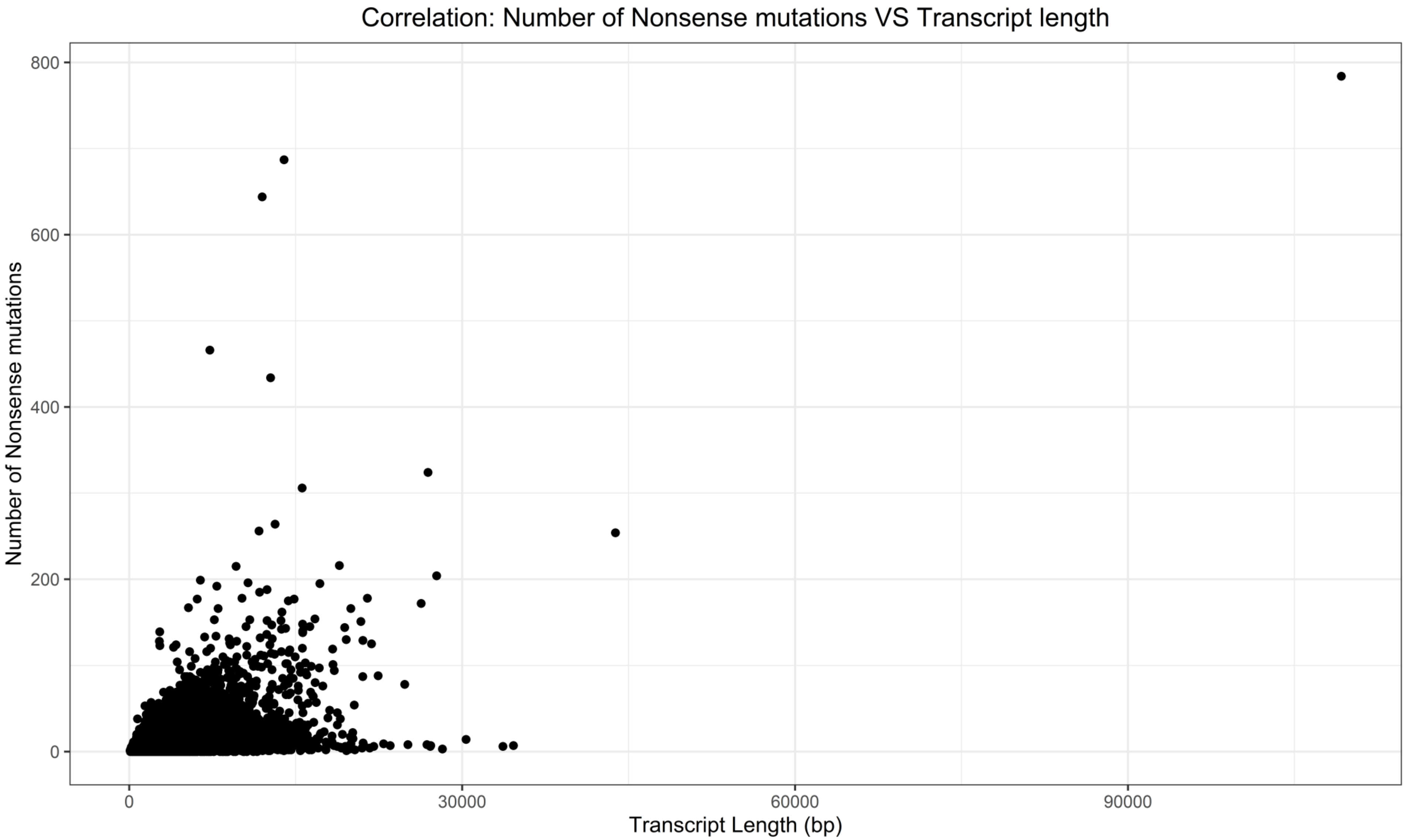


**Supplementary Figure 4J.**
Correlation between nonsense mutations and Transcript Length (bp) (Kendall test, tau = 0.21, p-value < 2.20E-16). The number of nonsense mutations and Transcript Length were obtained using biomart.

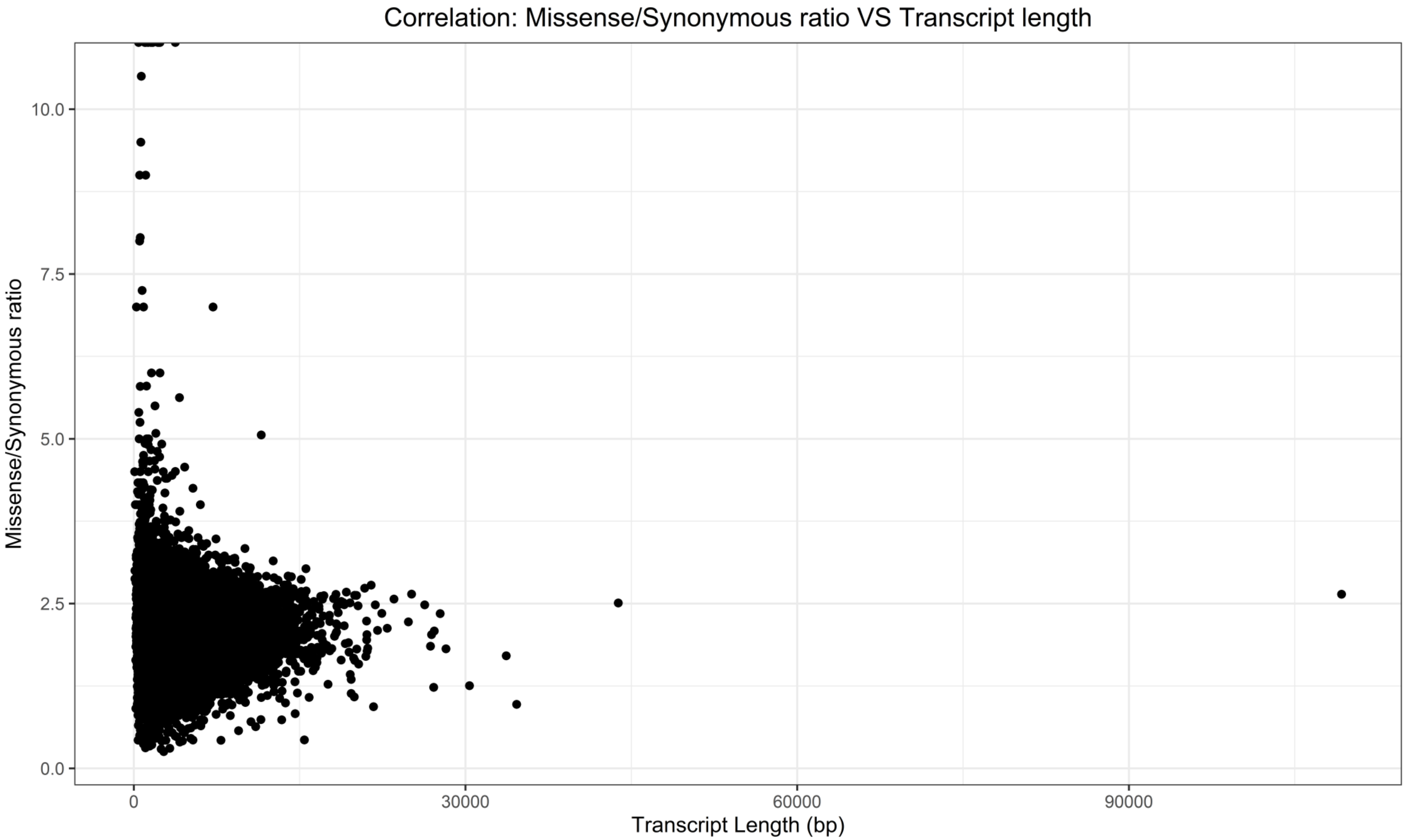


**Supplementary Figure 4K.**
Correlation between the missense/synonymous ratio of mutations and Transcript Length (bp) (Kendall test, tau = -0.07, p-value < 2.20E-16). The number of missense and synonymous mutations and Transcript Length were obtained using biomart.

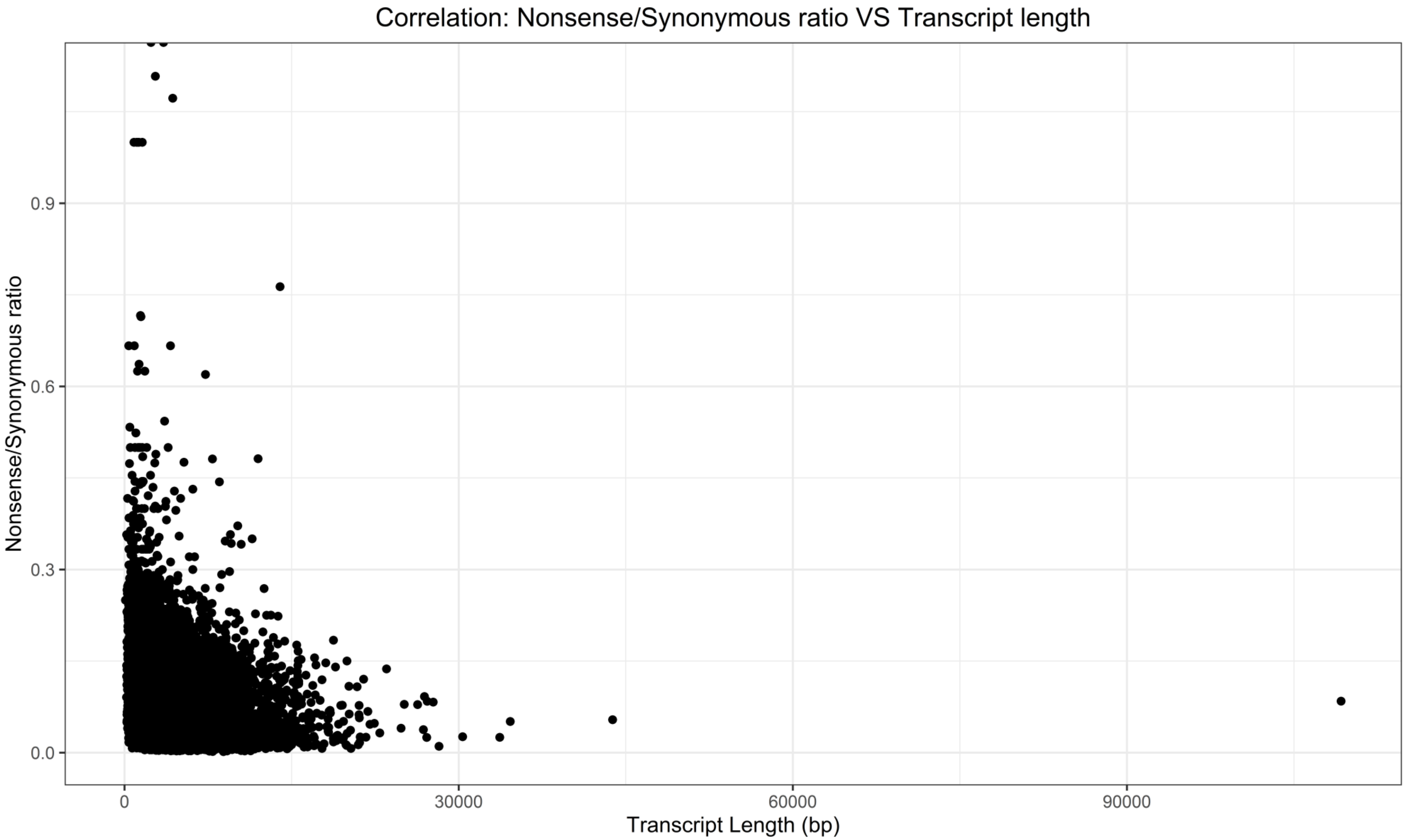


**Supplementary Figure 4L.**
Correlation between the nonsense/synonymous ratio of mutations and Transcript Length (bp) (Kendall test, tau = -0.19, p-value < 2.20E-16). The number of nonsense and synonymous mutations and Transcript Length were obtained using biomart.
