## Supplementary material for "Gene size matters: What determines gene length in the human genome?": S5 Fig

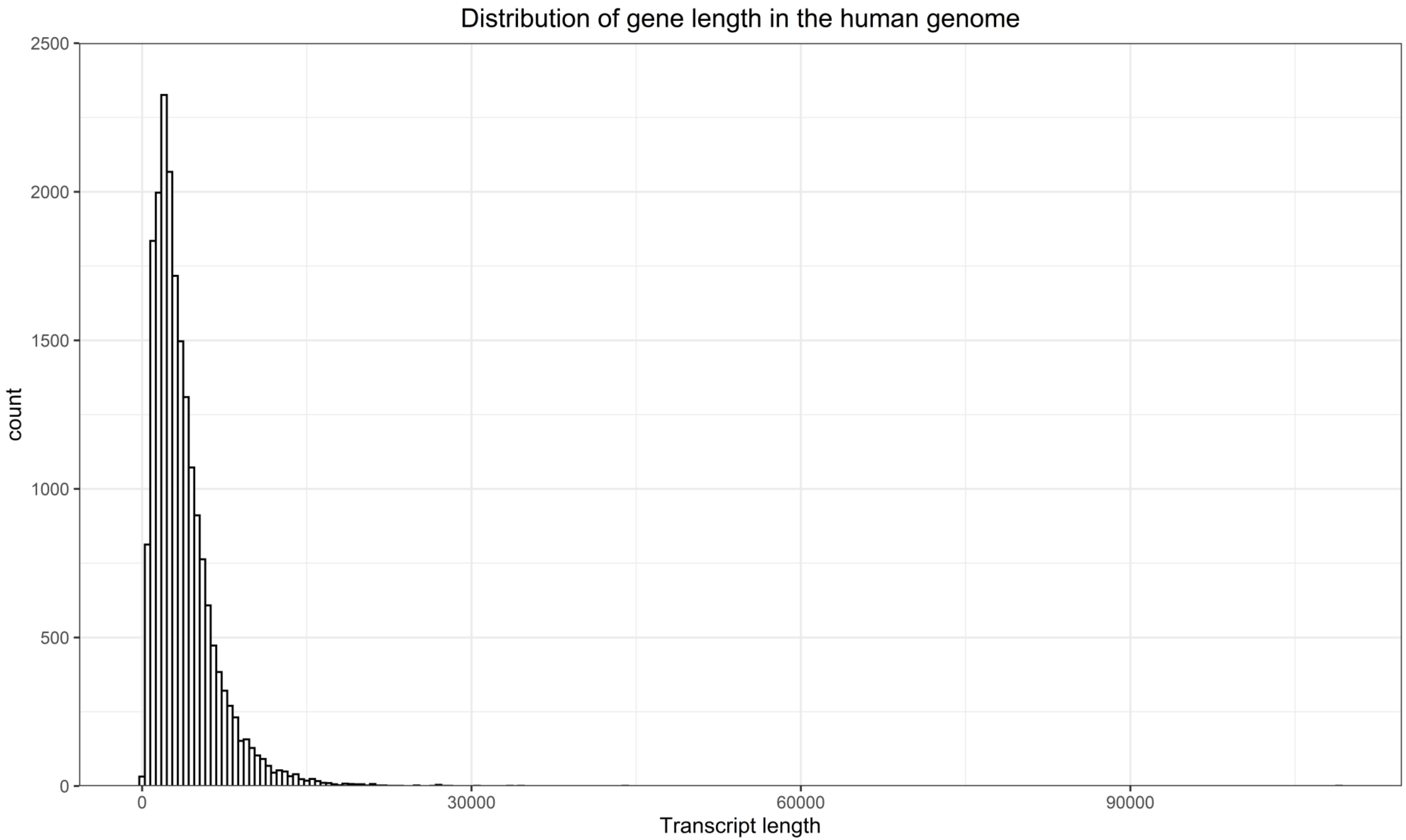


**Supplementary Figure 5A.**
Gene Length distribution in the human genome, for protein-coding genes only. Transcript Length was obtained from biomart.

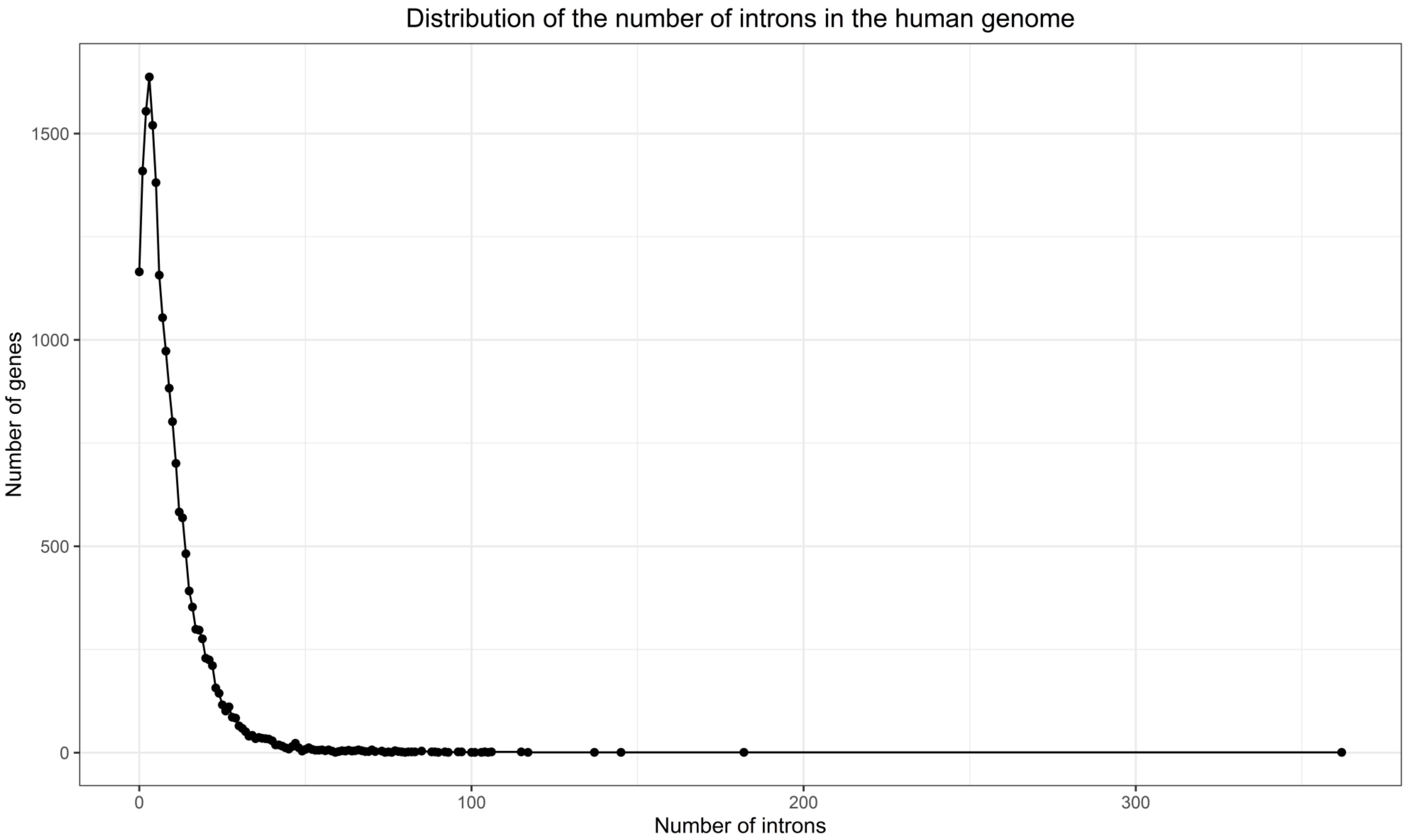


**Supplementary Figure 5B.**
Distributions of the number of introns in the human genome, for protein-coding genes only. The number of introns was obtained from biomart.
