## Supplementary material for "Gene size matters: What determines gene length in the human genome?": S6 Fig

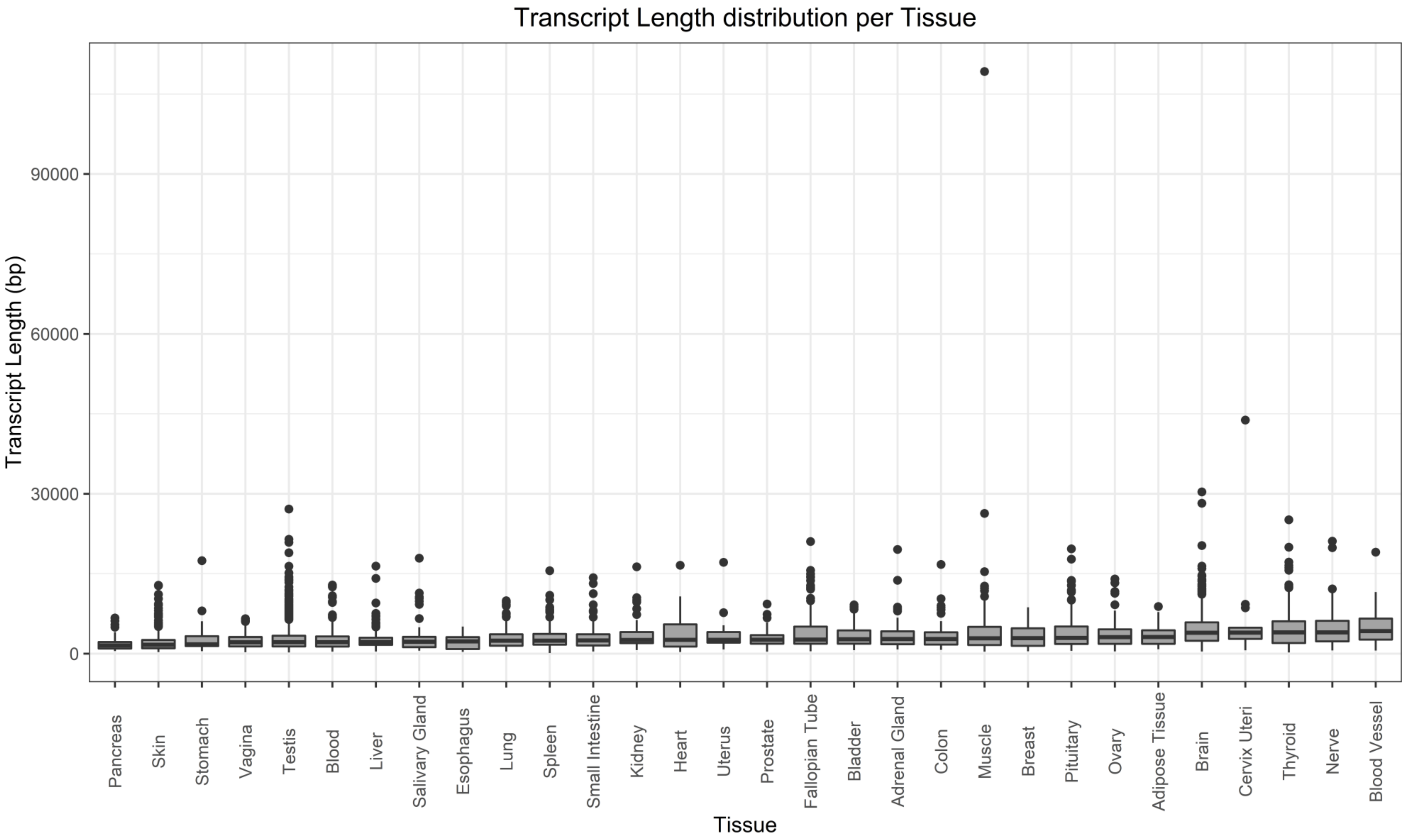


**Supplementary Figure 6.**
Transcript length distribution for genes specifically expressed in the given Tissues. Tissue specificity was defined as a gene having a Tau specificity score greater than 0.8.
