## Supplementary material for "Gene size matters: What determines gene length in the human genome?": S7 Fig

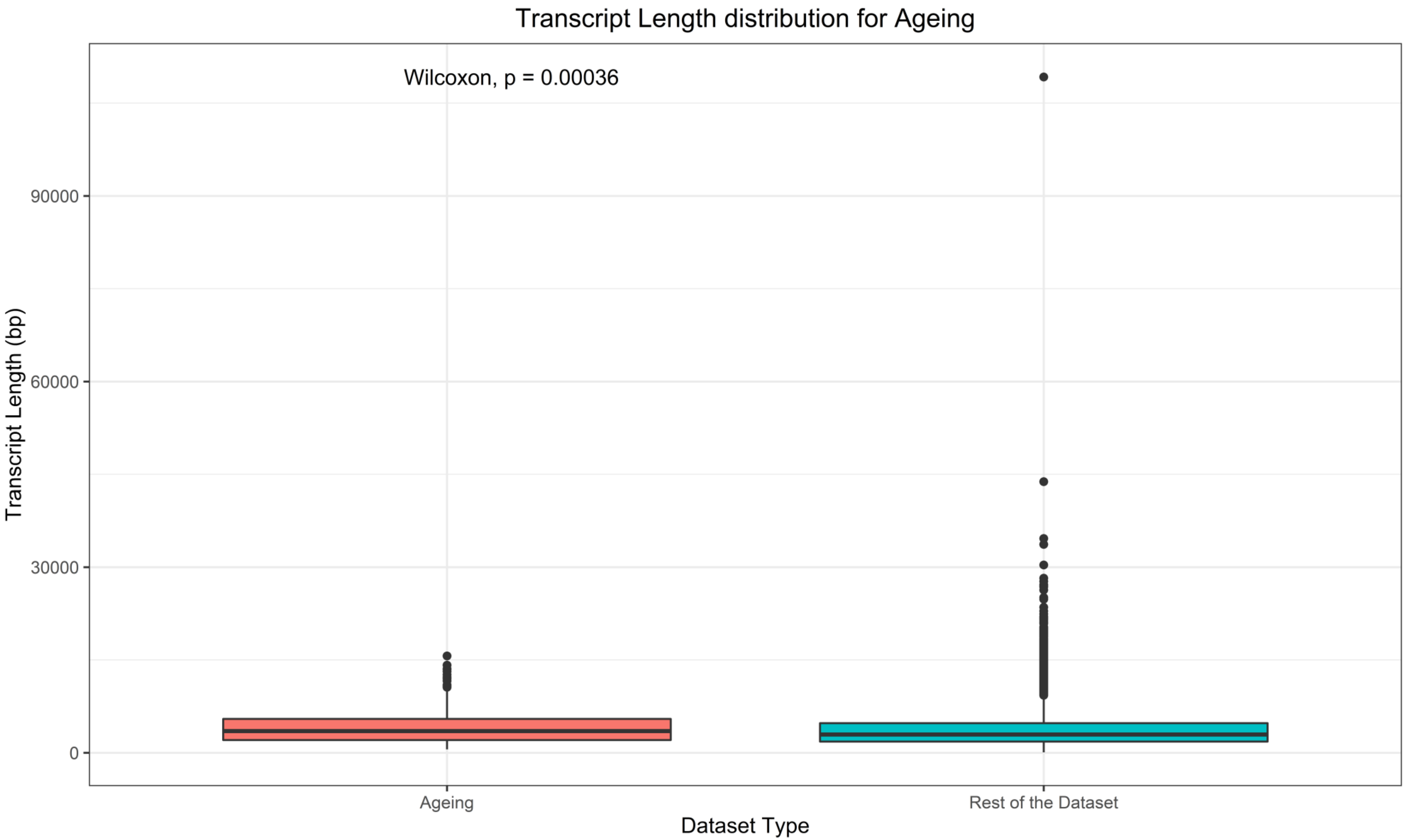


**Supplementary Figure 7A.**
Transcript length distribution for Ageing related genes and for the rest of the dataset. Ageing related genes were obtained from GenAge and Transcript Length was obtained from biomart.

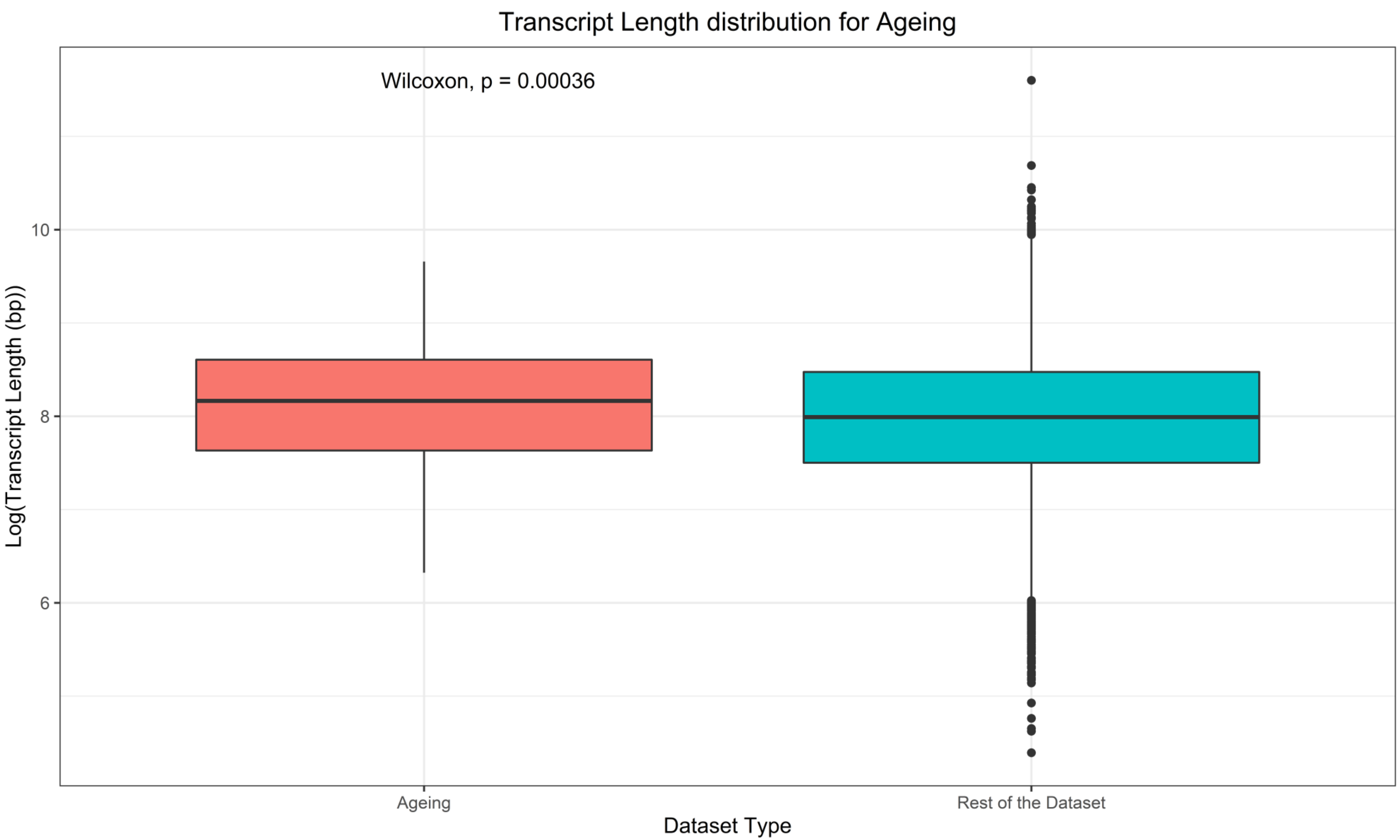


**Supplementary Figure 7B.**
Log transformed Transcript length distribution for Ageing related genes and for the rest of the dataset. Ageing related genes were obtained from GenAge and Transcript Length was obtained from biomart.

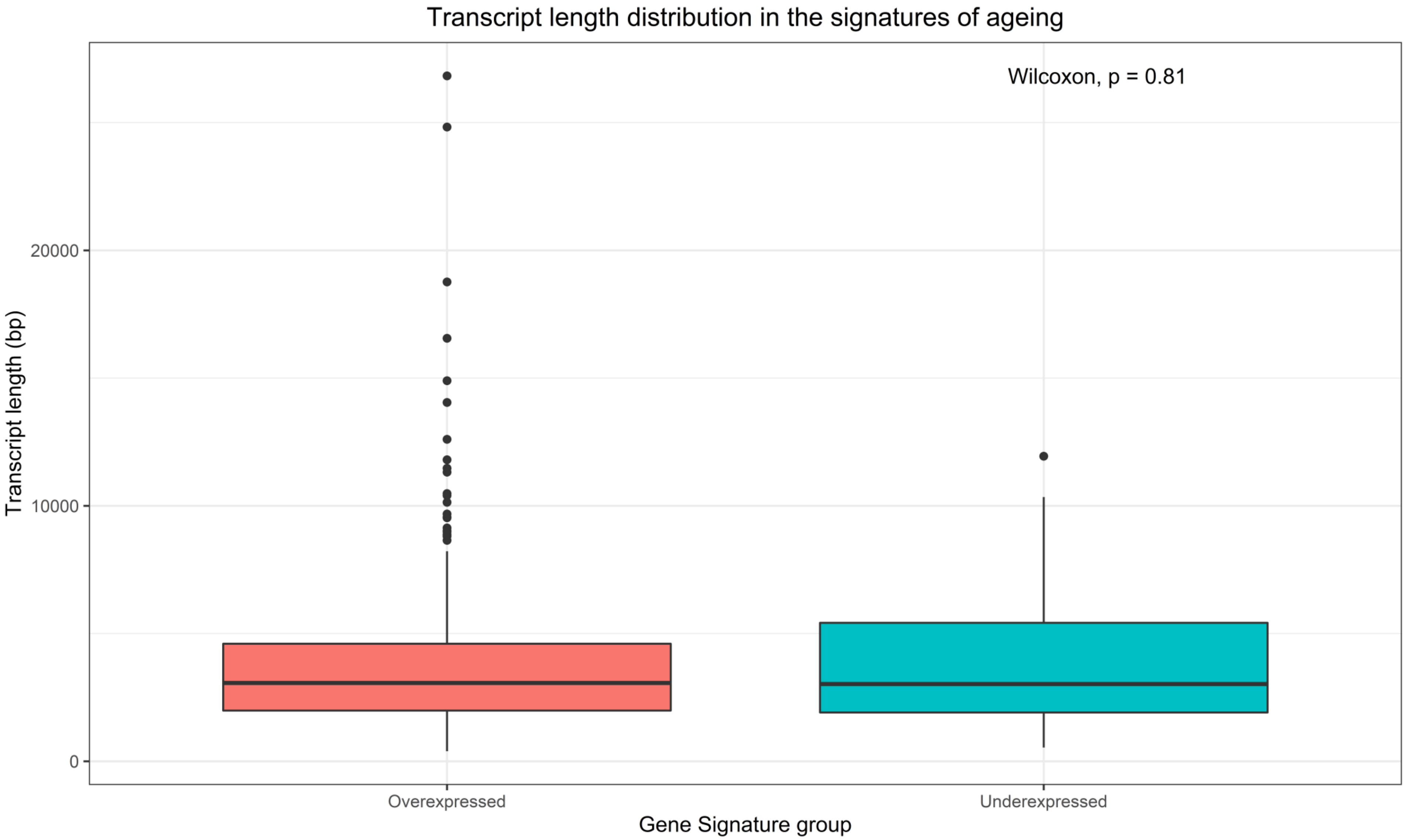


**Supplementary Figure 7C.**
Transcript length distribution for genes overexpressed and underexpressed with age.

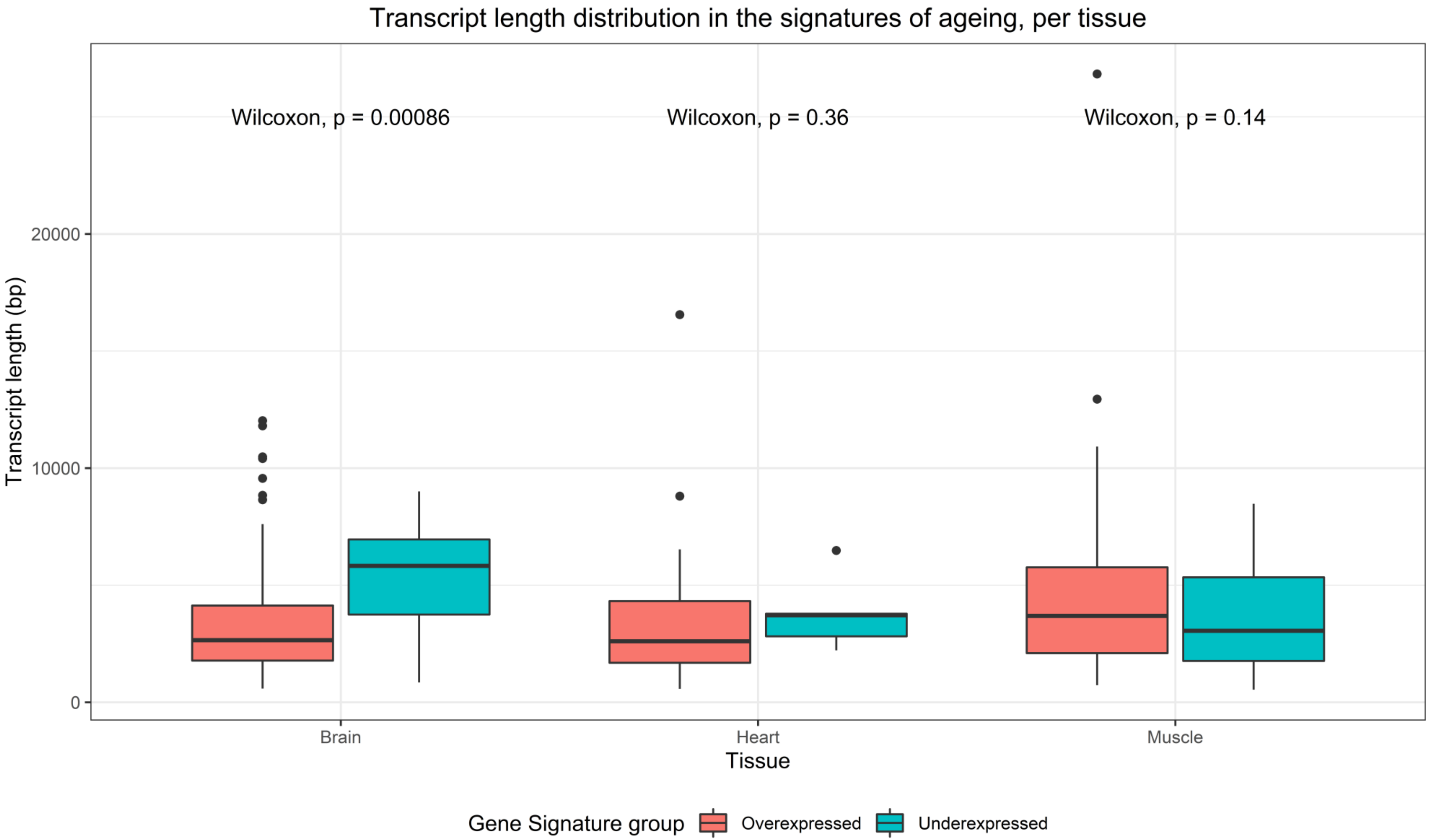


**Supplementary Figure 7D.**
Transcript length distribution for genes overexpressed and underexpressed with age, for brain, heart and muscle.
