## Supplementary material for "Gene size matters: What determines gene length in the human genome?": S8 Fig

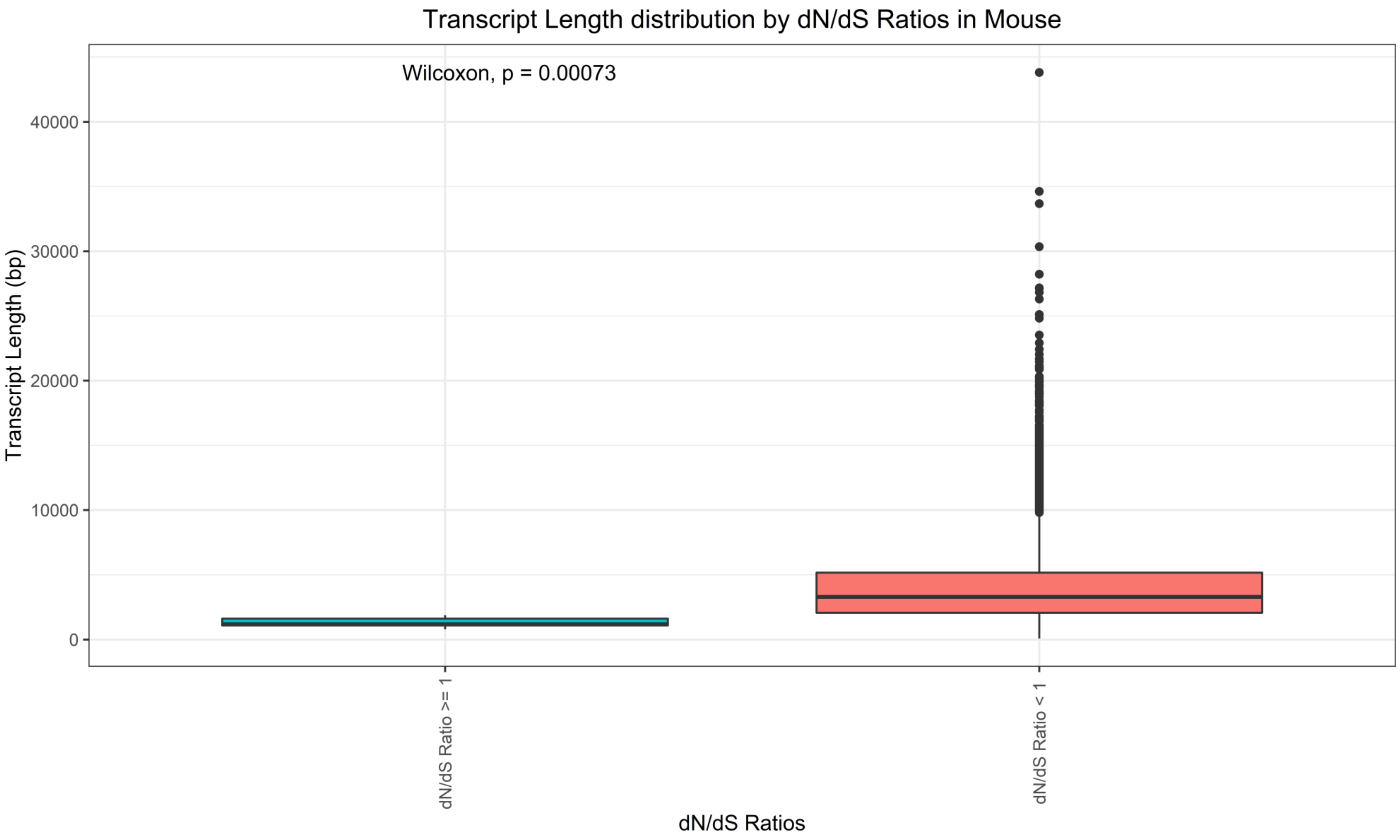


**Supplementary Figure 8A.**
Transcript length distribution for different dN/dS rations in Mouse. dN values, dS values and Transcript Length for each transcript were obtained using biomart.

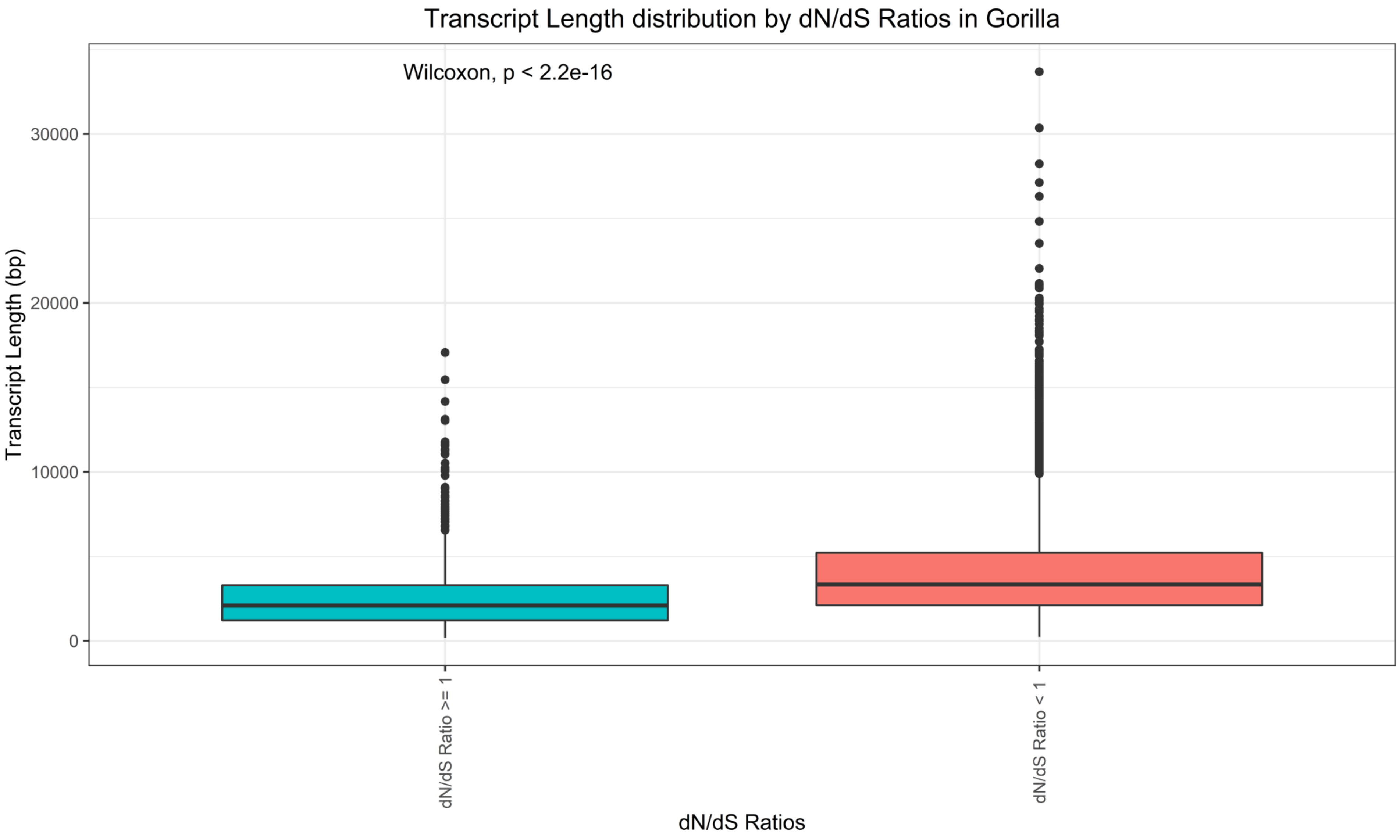


**Supplementary Figure 8B.**
Transcript length distribution for different dN/dS rations in Gorilla. dN values, dS values and Transcript Length for each transcript were obtained using biomart.

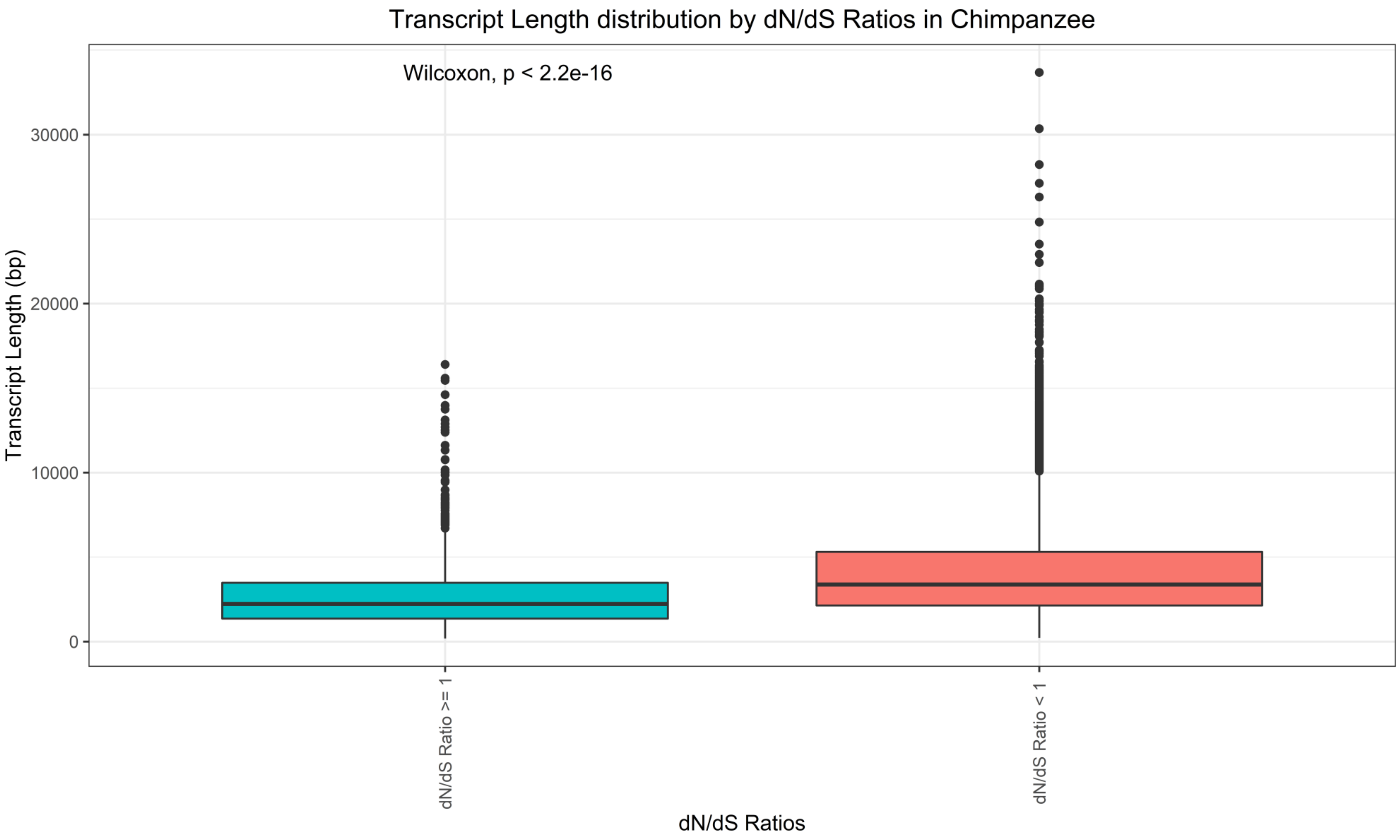


**Supplementary Figure 8C.**
Transcript length distribution for different dN/dS rations in Chimpanzee. dN values, dS values and Transcript Length for each transcript were obtained using biomart.
