## Supplementary material for "Gene size matters: What determines gene length in the human genome?": S9 Fig

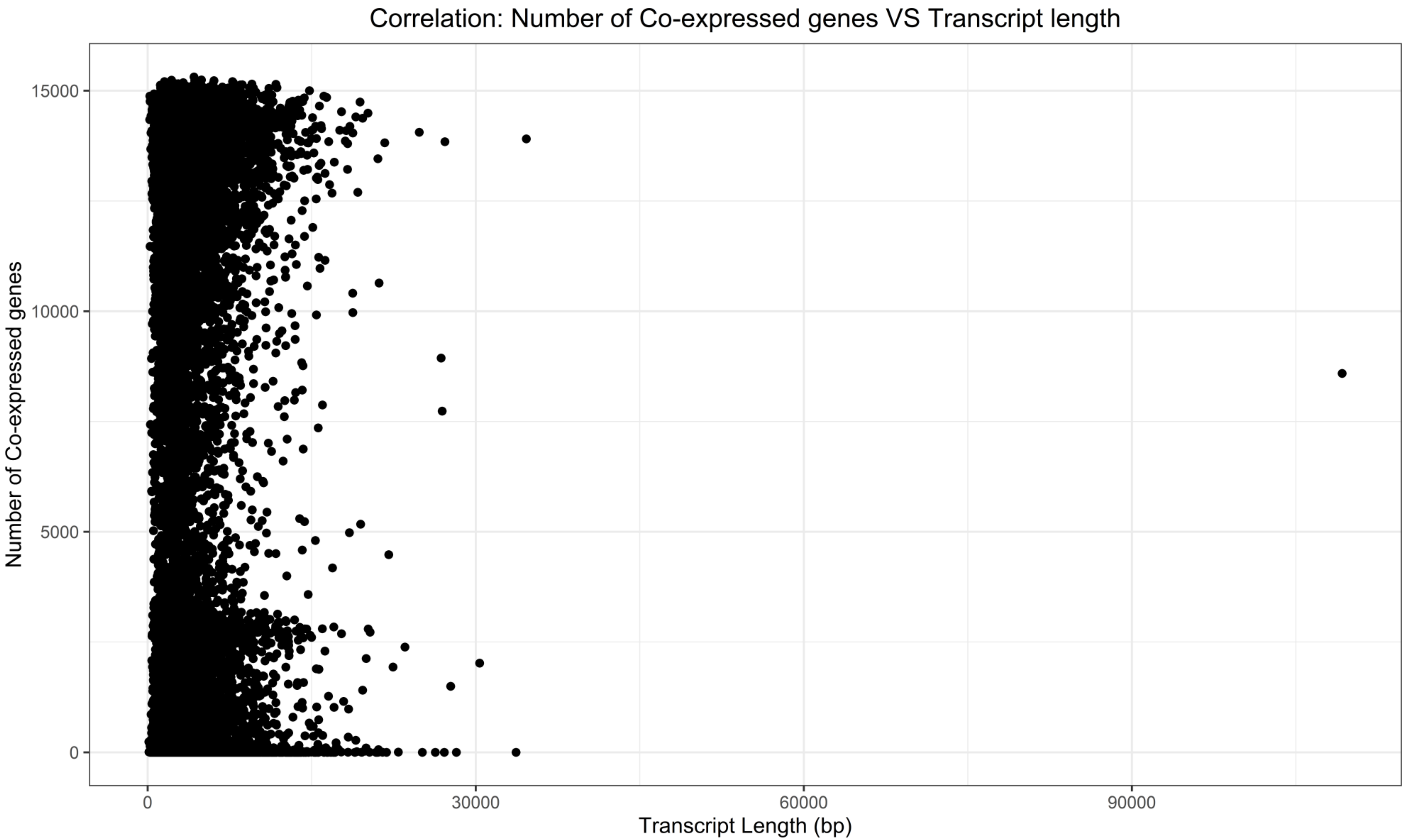


**Supplementary Figure 9A.**
Correlation between the number of co-expressed genes and Transcript Length (bp) (Kendall test, tau = 0.10, p-value < 2.20E-16). Number of co-expressed genes was obtained from GeneFriends and Transcript Length was obtained from biomart.

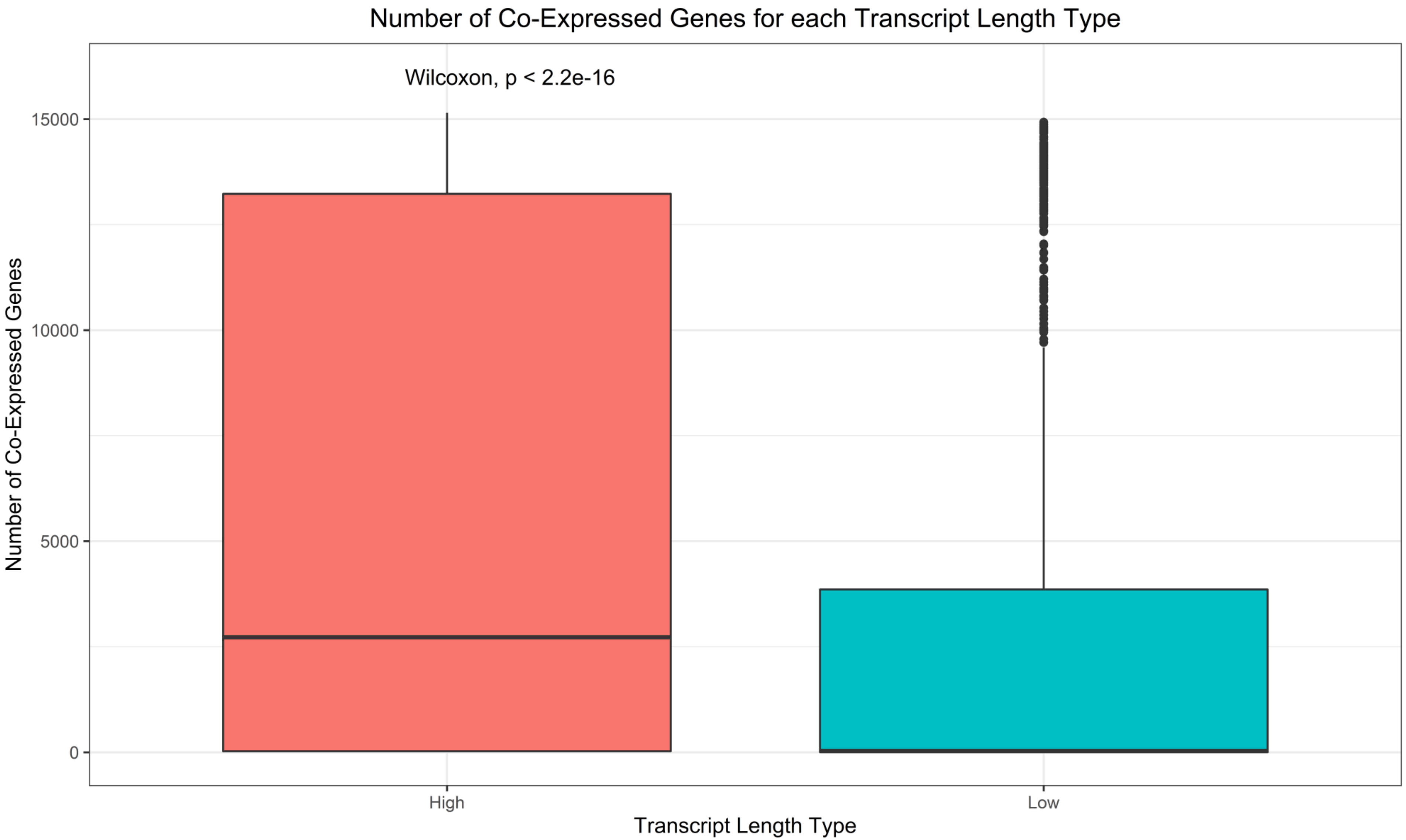


**Supplementary Figure 9B.**

Distribution of the number of co-expressed genes for long (High) genes and small (Low) genes. Number of co-expressed genes was obtained from GeneFriends and Transcript Length was obtained from biomart.

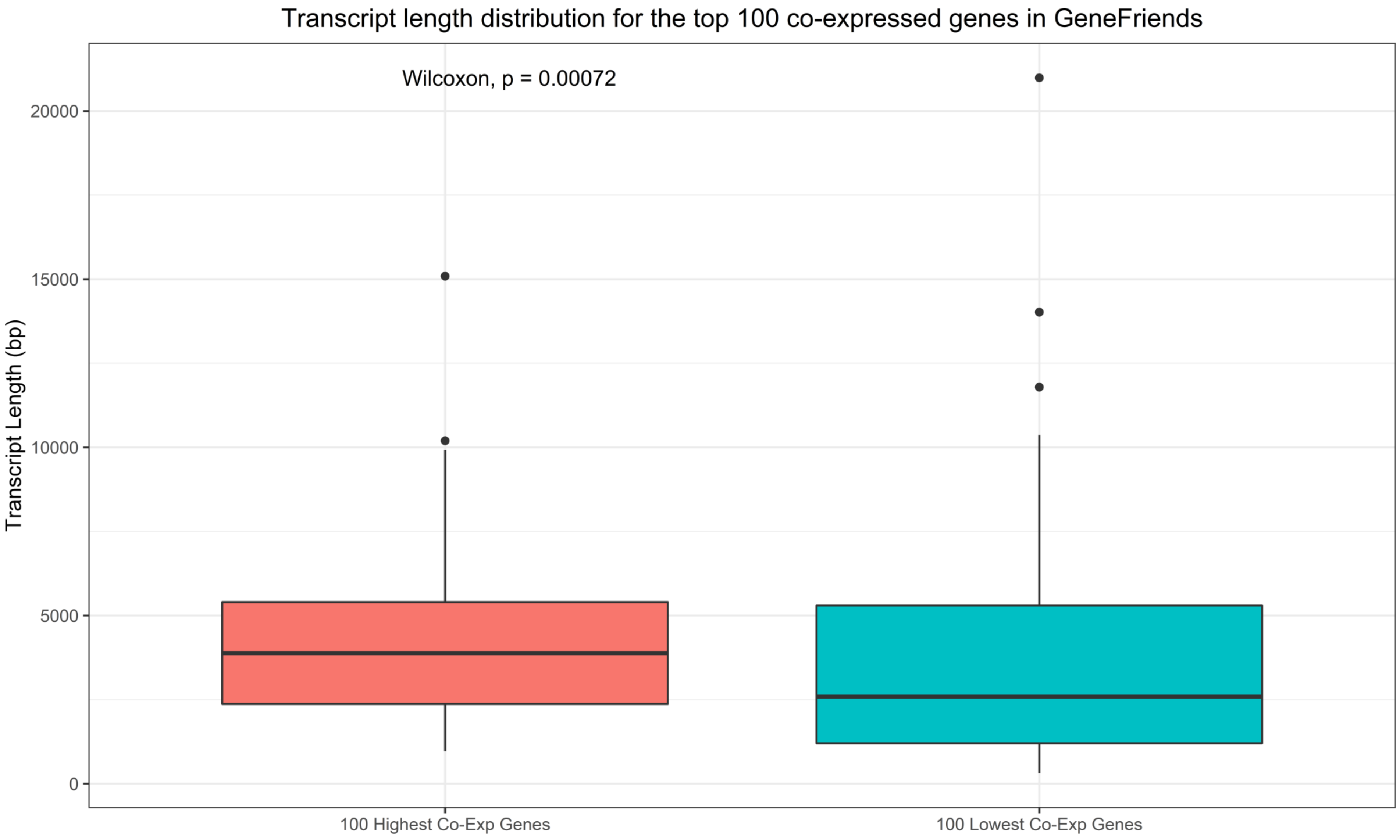


**Supplementary Figure 9C.**
Distribution of transcript length from our dataset for the top hundred highest/lowest co-expressed genes in GeneFriends. Median co-expression correlation values were calculated using GeneFriends and Transcript Length was obtained from biomart.
