## Supplementary material for "Gene size matters: What determines gene length in the human genome?": S10 Fig

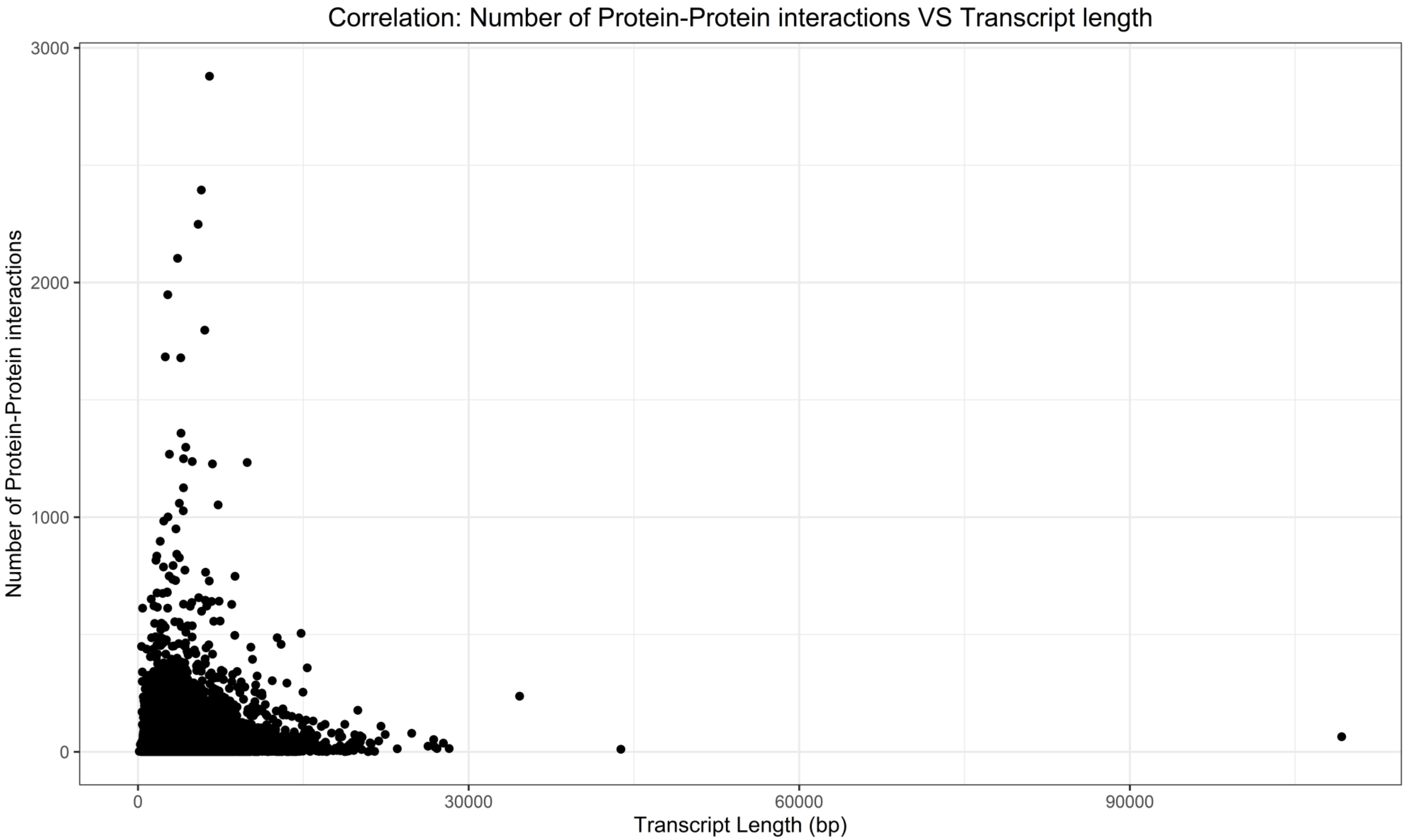


**Supplementary Figure 10A.**
Correlation between Number of Protein-Protein interactions and Transcript Length (bp) (Kendall test, tau = 0.06, p-value < 2.2E-16). Number of Protein-Protein interactions was obtained from BioGRID and the transcript length was obtained from biomart.

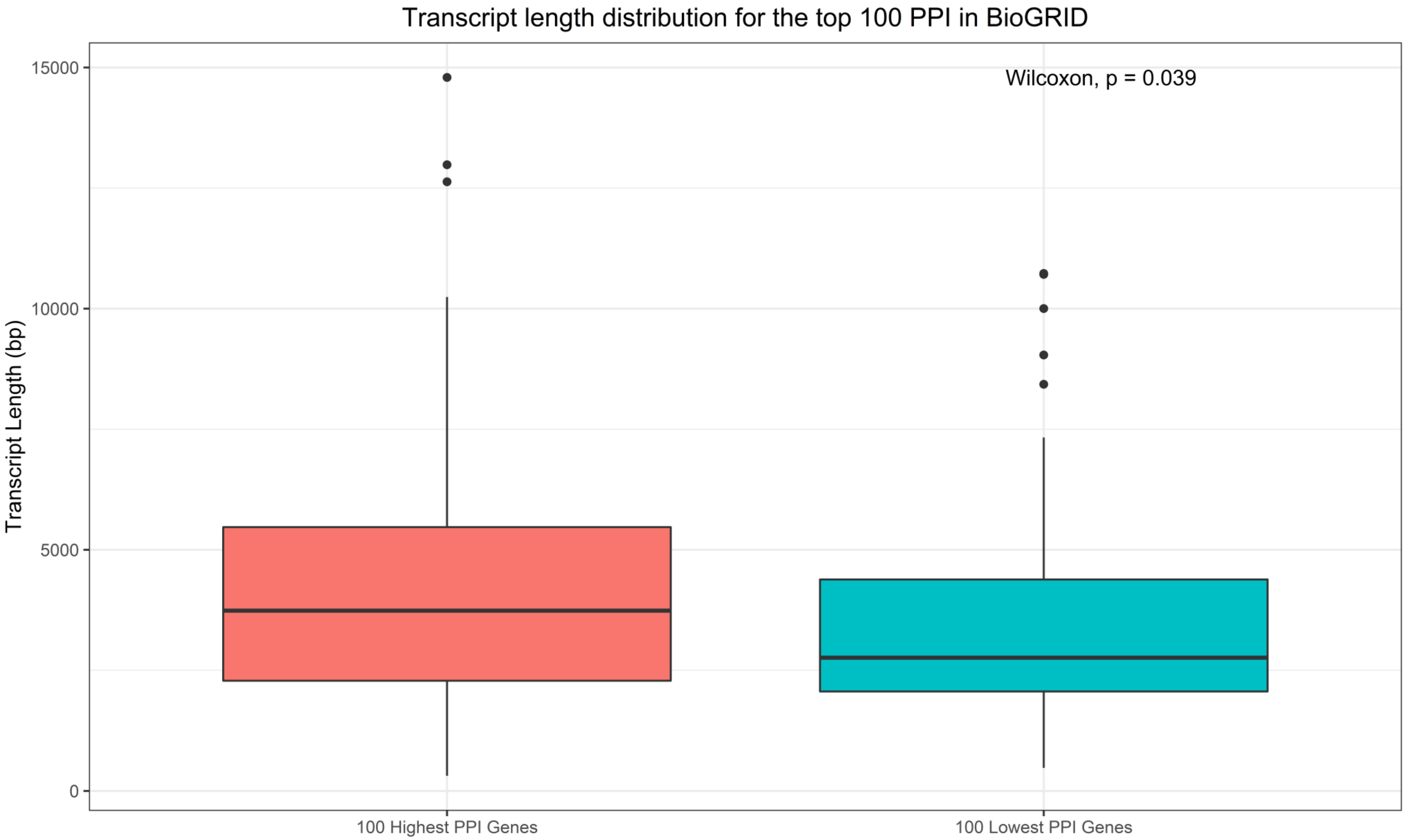


**Supplementary Figure 10B.**
Distribution of the transcript length for the top 100 genes with the highest and lowest protein-protein interactions. Number of Protein-Protein interactions was obtained from BioGRID and the transcript length was obtained from biomart.
